# Structural Modeling of *γ*-Secretase A*β_n_* Complex Formation and Substrate Processing

**DOI:** 10.1101/500488

**Authors:** M. Hitzenberger, M. Zacharias

## Abstract

The intra-membrane aspartyl protease *γ*-secretase (GSEC) cleaves single-span transmembrane helices including the C-terminal fragment of the amyloid precursor protein (APP). This substrate is initially cleaved at the ɛ-site followed by successive processing (trimming) events mostly in steps of three amino acids. GSEC is responsible for the formation of *N*-terminal APP amyloid-*β* (A *β*) peptides of different length (e.g. A*β*_42_) that can form aggregates involved in Alzheimer’s disease pathogenesis. The molecular mechanism of GSEC-APP substrate recognition is key for understanding how different peptide products are formed and could help in designing APP-selective modulators. Based on the known structure of apo GSEC and the APP-C99 fragment we have generated putative structural models of the initial binding in three different possible modes using extensive Molecular Dynamics (MD) simulations. The binding mode with the substrate helix located in a cleft between the transmembrane helices 2 and 3 of the presenilin subunit was identified as a most likely binding mode. Based on this arrangement the processing steps were investigated using restraint MD simulations to pull the scissile bond (for each processing step) into a transition like (cleavable) state. This allowed us to analyze in detail the motions and energetic contributions of participating residues. The structural model agrees qualitatively well with the influence of many mutations in GSEC and C99. It also explains the effects of inhibitors, cross-linking as well as spectroscopic data on GSEC substrate binding and can serve as working model for the future planning of structural and biochemical studies.

## INTRODUCTION

The intra-membrane aspartyl protease *γ*-secretase (GSEC) is a hetero-tetramer, consisting of the proteins nicastrin (NCT), presenilin (PS), anterior pharynx defective-1 (APH-1) and presenilin enhancer-2 (PEN-2).^1–4^ Presenilin with its nine transmembrane domains (TMDs), forms the catalytically active subunit of GSEC and contains the two active site aspartate residues, D257 and D385.^5^ Nicastrin, with 709 amino acids (AA), the largest of the GSEC proteins consists of one TMD and a large soluble ectodomain (ECD). The majority of the available literature attributes two main functions to NCT: Stabilizing the GSEC complex^6^ and preventing the inadvertent cleaving of non-substrates by impeding formation of the enyzme substrate (E-S) complex, due to steric clashes between the ECDs of NCT and the (non)-substrate.^1–3^ The roles of the two other proteins,^7^ APH-1 and PEN-2 involve the maturation of PS^4^ (PEN-2) and scaffolding^8^ (APH-1).

GSEC cleaves single-span transmembrane helices and so far more than 90 GSEC substrates have been uncovered.^1^ The most prominent of these substrates is C99, the C-terminal fragment of the amyloid precursor protein (APP). C99 is generated by *β*-secretase mediated cleaving of the extracellular domain of APP.^9^ After this pre-processing step, C99 can get into contact with GSEC and is then cleaved several times: At first C99 is reduced to a 49 (or 48) amino acid long fragment by an endo-peptidase step (also termed ɛ-cleavage), cleaving away the “amyloid precursor protein intracellular C-terminal domain” (AICD). Next, a succession of exo-peptidase cuts (*ζ*– and *γ*-cleavage) yields several mostly trimeric peptide fragments (from the C-terminus) as well as N-terminal amyloid-*β* (A*β*) peptides of different length.^10^ The main A*β* product is the 40 amino acid long A*β*_40_.^10, 11^ Several mutations and changes in temperature or membrane composition can shift this balance towards the production of amyloids that contain more amino acids^12,13^ – mainly A*β*_42_, but also even longer variants are obtained.

Although A*β*_40_ is known to form aggregates,^14^ the even more fibrillogenic A*β*_42_ has been identified to be the main component of the plaques found in post-mortem brain tissue of Alzheimer’s disease (AD) patients.^15,16^ These circumstances link both, C99 and GSEC closely to AD, the most common cause of dementia^17^ and puts them into the spotlight for the development of novel therapeutic strategies.

One possible approach is to prevent the production of overly aggregation prone (=longer) A*β* peptides in the first place. Thus, the main question regarding C99 processing concerns the mechanism(s) leading to the generation of such longer A*β* variants. Many experimental studies strongly suggest (1) that mutations at specific residues, either in GSEC or in C99 can influence the location of the (initial) *ϵ*-cleavage site^2,18–20^ and (2) that while processing as an exo-peptidase *γ*-secretase, cuts the A*β_n_* fragment preferentially in steps of three consecutive residues.^19,21,22^ This indicates that two different major A*β_n_* production lines exist: One starting from A*β*_49_ (*ϵ*-cleavage between C99 residues 49 and 50) and one where A*β*_48_ forms the endo-peptidase product (*ϵ*-cleavage between C99 residues 48 and 49). The first production line then leads in steps of three amino acids to A*β*_46_, A*β*_43_ and finally to A*β*_40_ which subsequently dissociates from the enzyme. The more pathogenic production line includes A*β*_45_ and A*β*_42_ where processing ends, releasing a longer end product into the extracellular matrix. Complimentary, the sequential processing model suggests that some of the known familial Alzheimer’s disease (FAD) – causing mutations lead to a destabilization of the A*β_n_*-GSEC complex which subsequently leads to the production of longer amyloids.^13,23^ In recent years, significant progress has been made in the investigation of *γ*-secretase. For instance, the CryoEM structure of GSEC (in different conformational states)^3,4^ has been solved and invaluable insights concerning the role of crucial C99 and GSEC mutations have been gained.^13,18,19,24–26^ However, important pieces of the puzzle surrounding A*β* production are still missing: Above all, the molecular mechanisms of how C99/PS mutations or GSEC modulators^23,27^ influence the outcome of C99 processing are not yet fully uncovered. This prevents rational, structure-based approaches to drug design and the development of new therapeutic strategies as well. Furthermore, the mechanism responsible for the repositioning of the A*β_n_* processing intermediates is not well understood.

Many of these issues could be addressed with a structure of the GSEC-C99 complex. Although not yet available, a number of studies have provided circumstantial evidence as to where the main GSEC-C99 interaction site might be located: Experimental investigations, such as photo-affinity mapping, have shown that C99 binds predominantly to the N-terminal domain of PS-1.^28^ Another very important hint has been given by a co-purifying helix in two of the reported CryoEM structures by Bai et al.:^4^ While the exact nature of the amino acid chain could not be elucidated, its position in a cavity, formed by TMDs 2 3 and 5 hints at a possible helix binding site in PS-1 (see also figure 1a). Other experimental studies, however, suggested that C99 may interface with TMDs 2, 6 and 9.^29^ Recent molecular dynamics (MD) simulations have revealed that the cavity formed by TMDs 2, 3 and 5, but also TMDs 2, 6 and 9 could be possible binding sites for hydrophobic species.^30^ Then again, other theoretical studies favored TMDs 2 and 6 as entry site^31^’^32^ for C99. All this data reduces the number of potential substrate entry (and binding) sites to three likely positions shown in Figure 1b.

**Figure 1:**
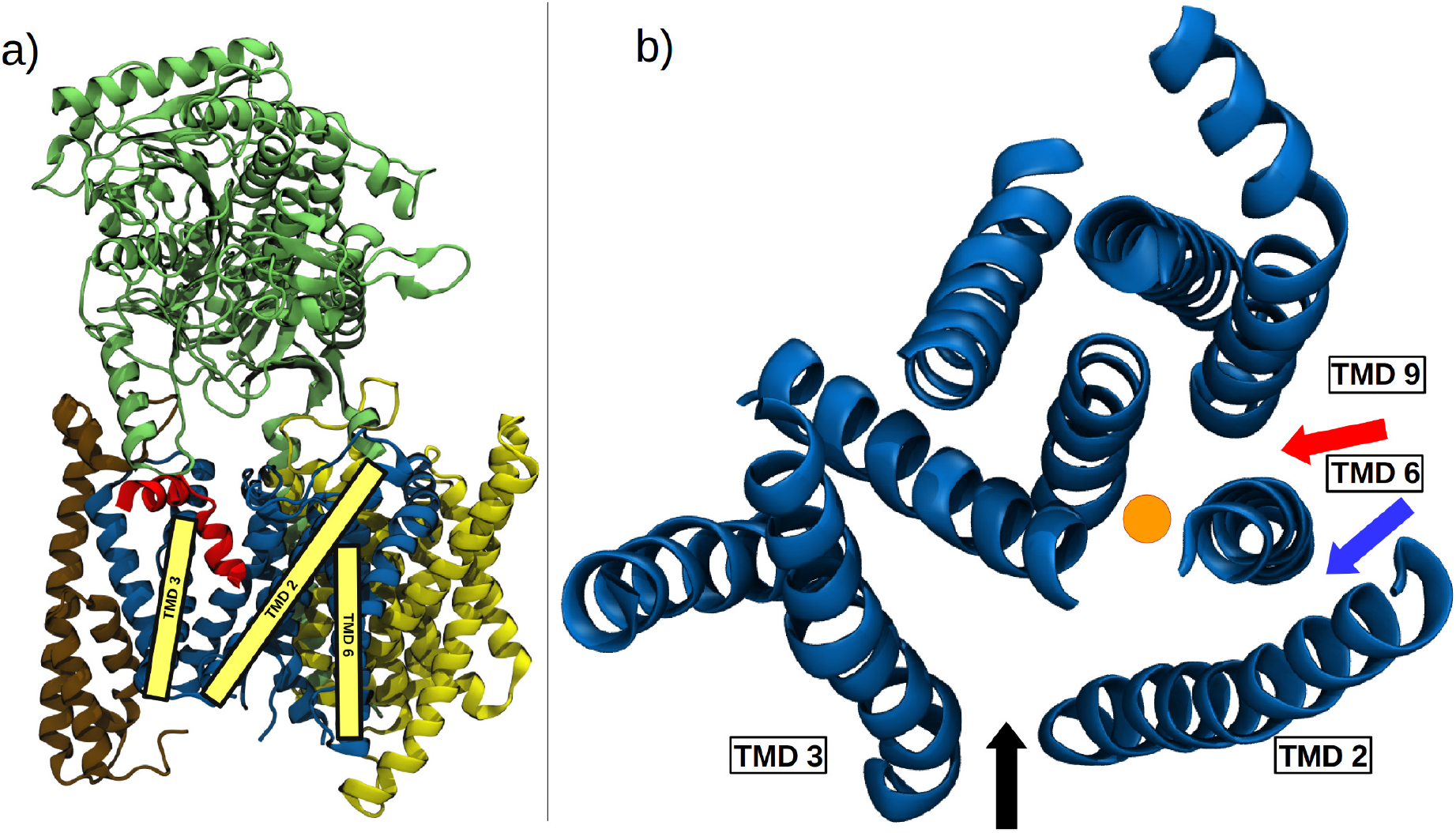
a) Cartoon representation of the structure of *γ*-secretase (PDB:5FN3^4^), consisting of nicastrin (NCT, green), presenilin (PS-1, blue), anterior pharynx defective-1 (APH-1, yellow) and presenilin enhancer-2 (PEN-2, brown). A co-purifying helix (red) bound in the cavity formed by TMDs 2, 3 and 5 is also shown. b) Three potential substrate entry routes towards the active site of PS-1 (blue cartoon, view perpendicular to the membrane plane, the approximate location of active site is indicated as yellow dot). In Model 1 (black arrow), the C99 substrate enters between TMD 2 and 3 and ends up in the cavity formed by TMDs 2, 3 and 5. The blue arrow depicts Model 2, while Model 3 where C99 interfaces with TMDs 6 and 9, is shown as a red arrow.

In the present study we performed atomistic *μ*s-timescale molecular dynamics simulations starting from the three possible GSEC-C99 complex geometries. By taking into account known structural data and through free energy calculations as well as interpretation of mutational and other experimental data we identified a most likely C99 binding site in the cavity formed by TMDs 2, 3 and 5. We then conducted *μ*s-timescale simulations of the putative GSEC-C99, A*β*_49_, A*β*_46_ and A*β*_43_ model complexes in order to profile key interaction sites and to gain insight into the mechanism of A*β_n_* processing. This allowed us to construct a working model of A*β_n_* processing, interpret the effect of mutations and to discuss an additional possible functional role of nicastrin.

## 1 RESULTS AND DISCUSSION

### 1.1 Identifying the main substrate binding interface

Putative models for the entry of C99 towards the active site of GSEC were generated by restraint MD simulations to initially bring the *ϵ*-cleavage site on C99 close to the active site, followed by extensive unrestraint simulations (see Methods for details). Simulations were started from three initial C99 placements resulting in models 1, 2 and 3 (Figure 2). However, in none of the simulations the *ϵ* sites remained in close contact with the catalytically active centers after removing the restraints: The centers of mass of L49/V50 (backbone) and D257/D385 (sidechain) were on average separated by 16.2 ± 0.7Å (Model 1), 9.9 ± 0.6Å (Model 2) and 15.9 ± 1.0Å (Model 3). This indicates that the investigated complexes represent resting states (with C99 positioned at exo-sites) occurring prior to the conformational change which guides the scissile bonds closer to the active site. However, the relatively large distance between the C-terminus of the substrate helix in Model 1 could explain why small transition state analogues (in contrast to inhibitory helices, see discussion in the final paragraph of the results section) cannot prevent the formation of the E-S complex:^33,34^ In the resting state, the N-terminus of C99 binds in the same cavity as a helical inhibitor is expected to and therefore both peptides would compete for the same binding site. The large space between the C-terminus of C99 and aspartates 257 and 385, however, could allow a small molecule inhibitor to access the active site without displacing the substrate, hence both molecules could bind at the same time.

**Figure 2:**
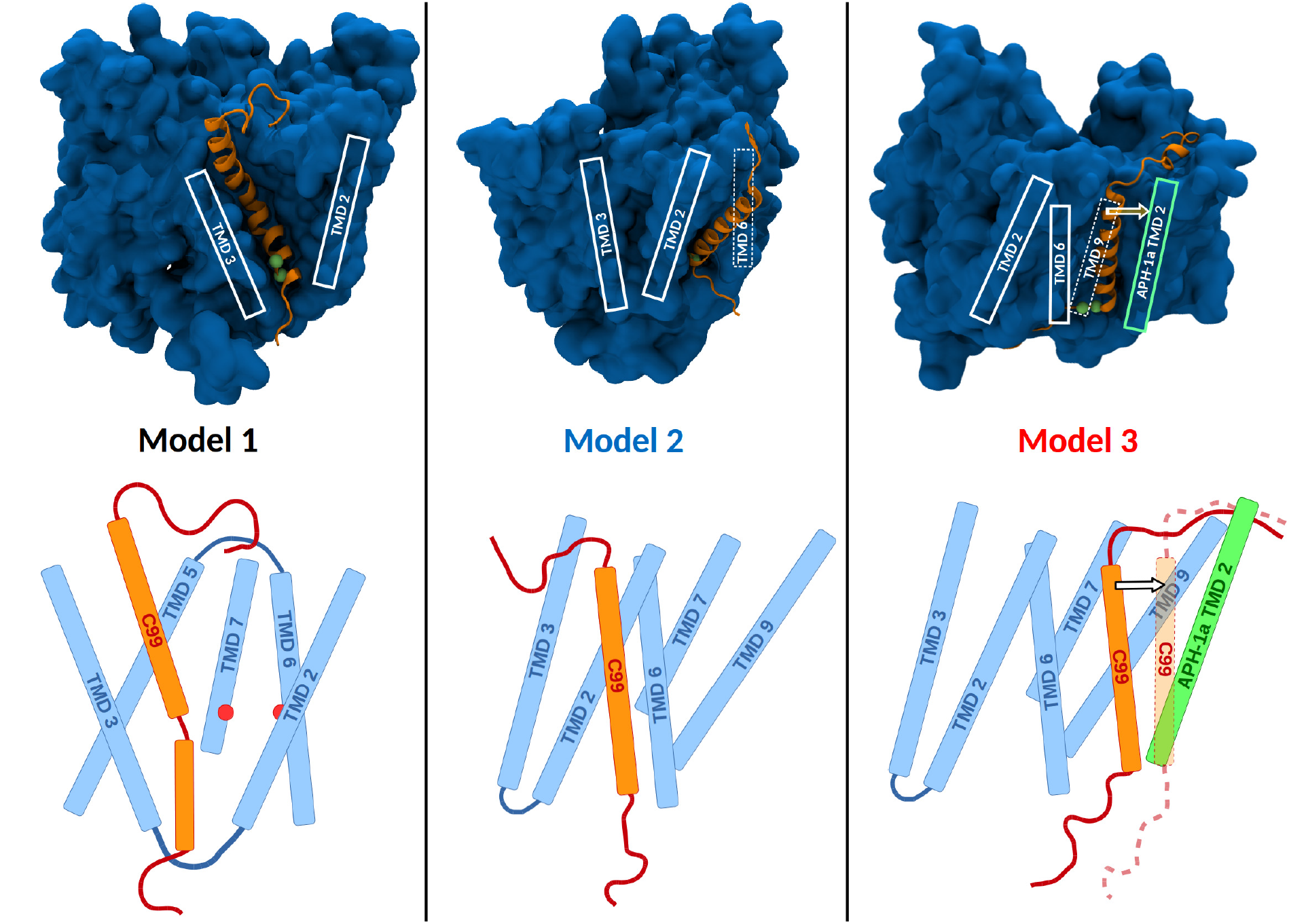
Depiction of the mean structures resulting from the Model 1 (left panel) Model 2 (center) and Model 3 (right) simulations. GSEC is depicted in blue, while C99 is orange. The *ϵ*-site is indicated by the C*α* carbons of residues L49 and L50 (green spheres). Bottom: Schematics representing the three models. The active site residues are indicated by red dots on the left panel.

Even though simulation trajectories of more than 3500ns have been generated, no conformational changes between the resting state and a putative transition state have been sampled. This suggests that such rearrangements are either very slow or very rare events and may be the reason for why C99 cleavage by GSEC occurs in time-scales of minutes.^1, 35^ Since it is impossible to produce simulation trajectories of sufficient length, we instead used steered MD to force the system into transition-like states for all investigations presented in the next sections.

In the simulations of models 1 and 2, the enzyme-substrate-complexes remained stable throughout the 3*μ*s sampling period. In the Model 3 trajectory, however, C99 was replaced by a POPC molecule at the TMD 6, 9 – interface around the 2*μ*s mark, while the substrate molecule moved away from the active site towards TMD 2 of APH-1a (see also right panel on Figure 2). To measure differences in mobility, all frames of each simulation have been aligned according to the C*α* atoms belonging to GSEC. From these aligned trajectories the mean B-factors of all C*α* atoms belonging to C99 residues G37 to V47 (this was the C99 region that adopted a helical secondary structure in all three simulations) have been calculated. The results confirmed the visual inspection of the trajectories and showed that C99 Model 3 displayed the highest mobility, with a mean B-factor of 156.0 Å^2^. Models 1 and 2 exhibited much lower mobilities, with Model 1 fluctuating less than Model 2 (51.3 vs. 68.7 Å^2^). Per-residue plots of the B-factor calculations can be found in the supplement.

From the obtained sampling trajectories of Models 1, 2 and 3, we also calculated the complex stability via MMPBSA and the values obtained clearly indicate that complex stabilization in Model 1, with -147.2 ± 0.5 kcal/mol is more favorable compared to models 2 (−97.7 ± 0.5 kcal/mol) and 3 (−108.2 ± 0.4kcal/mol). Please note that the large complex stabilization energies given in this section are due to the fact that entropy changes have been accounted for in the solvation model but not for complex formation, therefore the presented energies are closer to the binding enthalpy than the actual free energy (reasons for the omission of entropy changes are stated in the methods section).

C99 in Model 3 did not stay at the starting position and thus it can be argued that the general stabilization energy between C99 and the PS-surface lies at around 100kcal/mol (in absolute numbers). The complex stabilization energy of Model 2 is slightly lower than 100kcal/mol, which could be explained by the N-terminal loop region of C99: In models 1 and 3, the N-terminus binds steadily to either the loop 1 region (connecting TMD 1 with TMD 2) in case of Model 1 or the extracellular region of APH-1a (Model 3). Model 2, however, seems to prevent the formation of stable N-terminal C99-PS interactions, thereby decreasing the total complex stabilization. This is illustrated by the depiction of Model 2 in figure 2, where the N-terminus of C99 is not well defined (due to the structures depicting mean positions of all atoms in the respective simulation).

This indicates that TMDs 2 and 6 immobilize potential substrate molecules primarily via the formation of a small cleft in which the C-terminus of C99 can bind (see also the B-factor plot in the supplement, where the mobility of the C-terminal residues is greatly reduced in Model This putative binding pocket in Model 2 is relatively small and located close to the *ϵ*-site of C99 (see also middle panel on figure 2).

Interestingly, the mean solvent accessible surface area (SASA) is smaller in Model 3 (~ 3600Å^2^) than it is in Model 2 (~ 4000Å^2^). Therefore, strong binding of the C-terminal region seems to be the main reason for the reduced mobility of C99 in Model 2. The cavity formed by TMDs 2, 3 and 5 in Model 1 on the other hand, combines strong molecular attraction with steric trapping of a trans-membrane helix and a comparably small SASA of ~ 2800Å^2^ (indicating a higher degree of substrate burial).

Over 200 FAD causing mutations, affecting more than 130 different residues, have been reported on www.alzforum.org and it is likely that at least some of these directly steer E-S complex formation, stability or the positioning of the *ϵ*-site. In order to assess the potential of each model to explain the mode of action of these mutations, we counted the mean number of mutation sites that were within 5 Å of C99 in each system. With on average 23 residues, Model 1 is in direct contact with a way higher number of mutation sites than either Model 2 (average of 12) or Model 3 (average of 11), indicating that the substrate binding mode in Model 1 may be more susceptible to mutations and thus could better explain experimental findings.

It has been well established that GSEC exists in at least two functional states:^4^ active and inactive. Recent MD investigations indicate that GSEC can freely switch between these two conformations.^30, 36^ The two states are characterized by the distance between the active site aspartates, D257 and D385. As can be seen on table 1, presenilin in the Model 1 simulation is in a catalytically active state in a majority of sampled frames, while models 2 and 3 occupy the inactive state most of the time. Even though this finding can be due to non-exhaustive sampling of the phase space of each system, it would make sense that the binding of a substrate molecule at the active site leads to a higher probability for an enzyme to occupy the active conformation. The presence of a co-purifying helix in some of the CryoEM structures published by Bai et al.^4^ is further evidence that Model 1 represents a likely C99-GSEC complex.

**Table 1:** Mean distances between the PS active site aspartates D257 and D385. Values for C*α* and C *γ* distances are given.

| Mean distance of D257-D385 | Apo active<sup>†</sup> | Apo inactive<sup>†</sup> | Model 1 | Model 2 | Model 3 |
| --- | --- | --- | --- | --- | --- |
| Cα-Cα distance in Å | 8.3 | 9.1 | 8.2 | 9.6 | 10.0 |
| Cγ-Cγ distance in Å | 6.0 | 7.9 | 5.4 | 7.6 | 8.0 |

<sup>†</sup> Distances found in previously published Apo-structure simulations<sup>30</sup> have been added for comparison.

Two arguments speaking against TMDs 2 and 3 forming the entry to the active site in the TMD 2, 3, 5 – cavity are (1) the short loop connecting TMDs 2 and 3 and (2) the shorter *ϵ*-site distances in the Model 2 simulation, indicating that the conformational change leading to the transition state would have to be less pronounced in the Model 2 case. The first argument, invokes two requirements for complex formation of (uncleaved) C99 with GSEC: Firstly, the substrate, since it is too long to fit into GSEC in its entirety, has to be bent in order to allow the scissile bond to get into contact with the active site. After the initial *ϵ*-cleavage removes the AICD, the remaining A*β_n_* peptide fits the cavity without bending. In the Model 1 simulation, C99 is kinked in the region between G37 and V40, an area often referred to as the “hinge region”.^2^ This requirement for helix bending may explain why *γ*-secretase substrate flexibility has very often been found to be connected to enzyme binding and proteolytic activity.^2,37–41^

The other requirement is that the distance between TMDs 2 and 3 increases, so that a protein helix can enter the active site. In the Model 1 simulation, TMD 2 moves away from TMD 3 as C99 is pulled inside the cavity. This fits the experimental^3, 4^ and theoretical findings^30,42^ attributing heightened mobility and flexibility to TMD 2 (including the N-terminus of TMD and confirms its role as substrate entry gate.

In case of the second argument one has to note that the shorter *ϵ*-D257/D385 distances of Model 2 are only caused by the fact that the binding interface is spatially closer to the active site than in the other models. Conformationally there is nothing to suggest that C99 is more “ready to be cleaved” in Model 2 than it is in Model 1. In Model 2 the scissile bond still has to traverse inside the PS “pore” to get into contact with aspartic acids 257 and 385.

However, the putatively easier access to the active site and less stable complex in Model could also be taken as a hint for the presence of two different active binding sites: One (Model 2) for the initial cleavage and a second (Model 1), where the *γ* and *ζ* processing steps take place. The binding cavity of Model 1 is not easily accessible from the TMD 2, 6 interface since the extracellular loop 1 (between TMDs 1 and 2) blocks the direct path. In order to migrate from the Model 2 site to the binding cavity, the substrate’s N-terminus would have to somehow move underneath loop1 without becoming entangled, which for steric reasons is very unlikely and thus also highly impractical. The only realistic entry path would be dissociation from the TMD 2, 6 interface and migration across the surface of PS 1 until the TMD 2, 3 interface is reached. Even though both pathways seem to be very improbable, we still think that at least the second pathway (migration across the enzyme surface) can not be completely ruled out and further (experimental) investigations are necessary to assess this possibility more closely.

For efficiency reasons, the Model 1, 2 and 3 simulations have been conducted in absence of the NCT ECD, however, a superposition of Model 1 with the complete GSEC structure indicated that if C99 binds at the Model 1 location then it is very likely also in contact with a small patch of the NCT ECD, ranging from S241 to E245. This contact region between NCT and the substrate coincides with the interface between a co-purified helix and nicastrin found in some of the CryoEM structures published by Bai et al.^4^ Therefore, the Model 1 binding mode would also explain best NCT-C99 interactions that have previously been suggested by experiment.^28^’^43^ Even though such interactions cannot be completely ruled out for Model 2, they are much more unlikely, due to the absence of a stable NCT-PS interface at TMDs 2 and 6. For all the reasons discussed above, in the subsequent parts of the manuscript we assume that Model 1 resembles most closely a A*β_n_*-GSEC complex with the *ϵ*-cleavage site in close proximity to the GSEC active site and discarded the two other hypotheses. We also included the nicastrin ectodomain because of the putative PS-NCT interface.

### 1.2 Nicastrin stabilizes C99 binding in Model 1

Since we simulated all GSEC-A*β_n_* complexes twice, once in presence and a second time in absence of the nicastrin ectodomain, we were able to compare complex stabilities (Table 2). The A*β_n_*-GSEC complexes were consistently more stable in the presence of nicastrin (mean AAG of ~ -30 kcal/mol). In simulations missing the NCT ECD, the N-term formed additional interactions with the loop 1 region. However, the MMBPSA calculations indicate that the stabilizing influence of the ECD outweighs the effect of additional PS-loop1-A*β_n_* interactions. Hence, it is likely that the role of NCT also lies in the stabilization of the E-S complex.

**Table 2:** Results of MMPBSA calculations of the A*β_n_*-GSEC complexes with and without the NCT ECD. Errors are given as standard errors of the mean (n=1500).

| Simulations | GC99(*) | GA$\beta_{49}$(*) | GA$\beta_{46}$(*) | GA$\beta_{43}$(*) |
| --- | --- | --- | --- | --- |
| ECD | -212.9 ±0.6 kcal/mol | -147.9 ±0.4 kcal/mol | -107.5 ±0.6 kcal/mol | -104.5 ±0.4 kcal/mol |
| no ECD | -154.9 ±0.8 kcal/mol | -125.1 ±0.4 kcal/mol | -91.9 ±0.5 kcal/mol | -80.8 ±0.7 kcal/mol |
| $\Delta\Delta G$ | -58.0 ±0.8 kcal/mol | -22.8 ±0.6 kcal/mol | -15.6 ±0.8 kcal/mol | -29.7 ±0.8 kcal/mol |

### 1.3 The substrate processing mechanism

In the GC99 simulation, the substrate ended up exactly at the same location as the co-purifying helix in the structures published by Bai et al.^4^ (a superposition of both structures can be found in the supplement).

In order to investigate further processing of the C99 substrate we generated model structures of GSEC in complex with A*β*_49_, A*β*_46_ and A*β*_43_ intermediates based on the Model 1 placement in the C99-GSEC complex. The models were obtained by restraint MD-simulations to move the expected cleavage site of the shorter substrates (scissile bond) into the vicinity of the active site (see Methods for details).

Generally, two different modes of action are possible: The helix could unwind and elongate so that the scissile bond can get into contact with the active site aspartate residues or the substrate could slide deeper into the binding cavity, thereby keeping its helical structure.

As can be taken from figure 3, the simulations showed that between each cleavage step, the substrate is pulled deeper into the binding cavity of PS. However, the data also suggests that there is an N-terminal anchoring region that remains in place while A*β_n_* is processed. Furthermore, the distance plot on figure 3 indicates that at the last processing step, from A*β*_43_ to A*β*_40_, the N-terminal anchor begins to shorten (destabilize): In simulations GA*β*_49_ and GA*β*_46_ the anchoring regions ranged from V12 (the first C99 residue in the model) until D23 (or even V24 if standard deviations are taken into account as well). In GA*β*_43_, the anchor was one residue shorter, as D23 was already pulled a little bit further inside the lipid membrane region (there is no overlap between the standard deviations of D23 in GA*β*_43_ and the other simulations), suggesting beginning destabilization of the anchoring region. Further destabilization of the N-terminal anchor as it would be the case in A*β*_42_ can be expected to lead to a shortened lifetime of the E-S complex and may be the reason why APP processing is more likely to end at this point. A similar N-terminal anchoring region, considered to bind to the N-terminal fragment of PSEN and its putative stabilizing effect has previously been proposed by Chávez-Gutiérrez et al. who also suggested that the substrate unwinds after it is cleaved by GSEC.^13^

**Figure 3:**
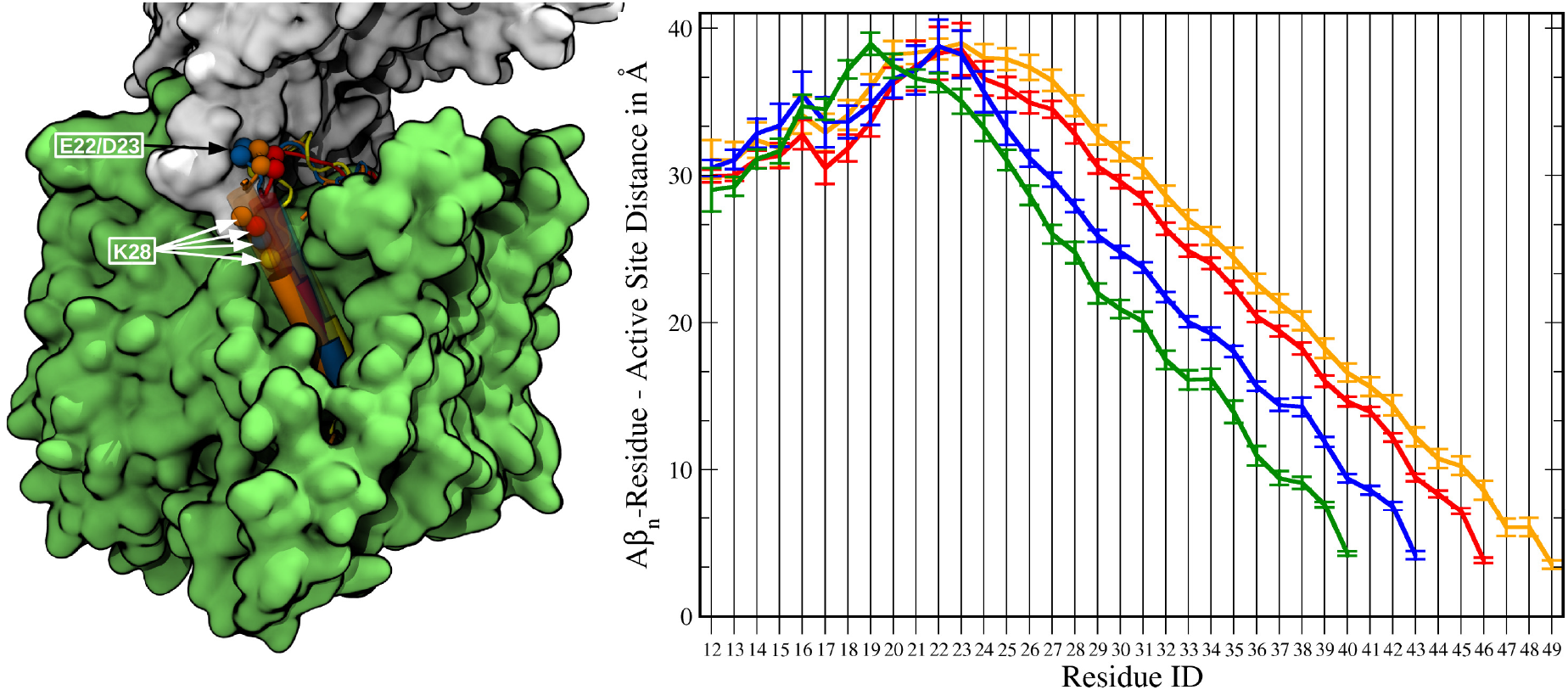
Left panel: GSEC (TMDs green, NCT ECD white) in complex with C99 (orange), A*β*_49_ (red), A*β*_46_ (blue) and A*β*_43_ (yellow). K28 travels deeper into the hydrophobic region, while the chain around E22/D23 remains in position. Right panel: Mean COM distance between the backbone of each residue and the active site side chains. The error bars represent the standard deviation. GC99 is depicted in orange, GA*β*_49_ in red, GA*β*_46_ in blue and GA*β*_43_ in green. The absence of an overlap of the error bars of residues (Residue ID >24) indicates a systematic displacement of the respective helix segment towards the active site.)

Analysis of the extend of helicity of the substrate molecules, measured via characteristic hydrogen-bonds, in our simulations advocate for a combined sliding/unwinding mechanism: In Figure 4a, residue n in frame i has been deemed helical if its backbone amide group either functioned as an H-bond donor to the n-4*th* residue or as an H-bond acceptor for the n+4*th* residue. A distance-based criterion has been used to determine the existence of H-bonds (distance between donor-H and acceptor O < 3Å).

**Figure 4:**
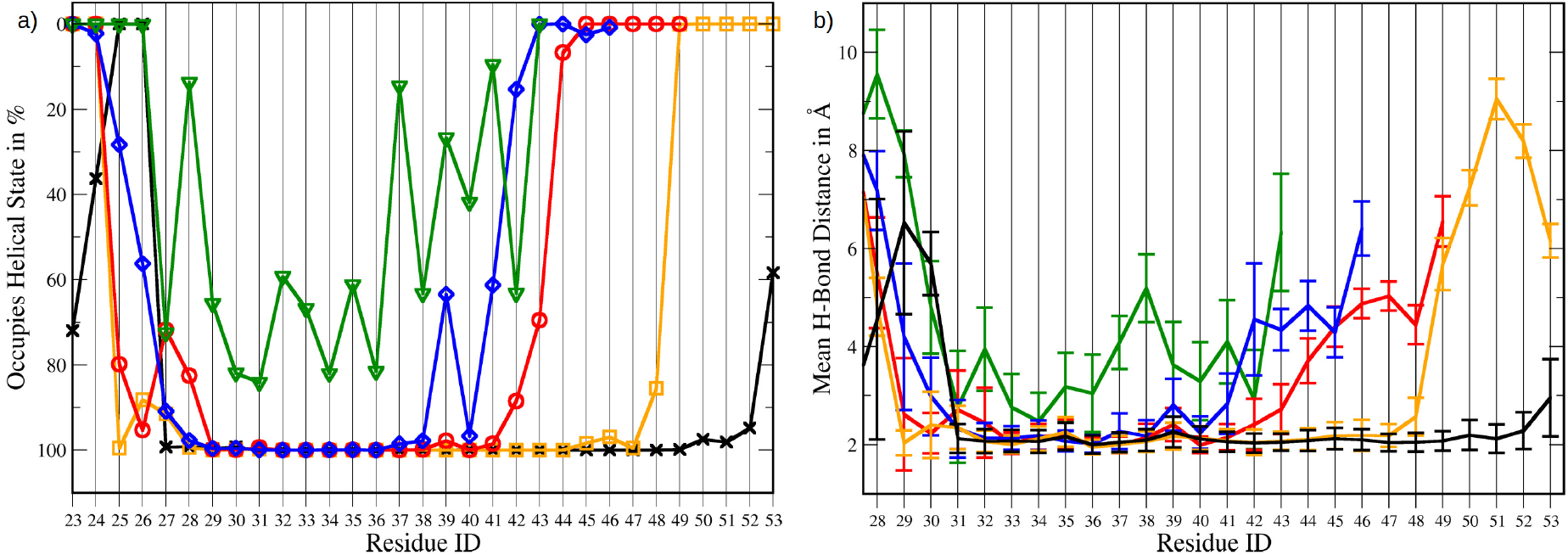
a) Occupation of the helical state in % of evaluated frames. The result for free C99 is plotted in black, GSEC bound C99 (simulation GC99) in orange, GA*β*_49_ in red, GA*β*_46_ in blue and GA*β*_43_ in green. b) Mean H-bond length between the n*th* the n-4*th* residue. The color scheme is the same as above and the error bars depict the standard deviation.

The helicity calculations clearly show that every processing intermediate is destabilized at the C-terminus and considerably less helical than its predecessor. Also, the N-terminus is unwinding during substrate processing which is necessary for the anchor region to stay in place while the remaining helix is moving further into the binding site. While C99, A*β*_49_ and A*β*_46_ remain relatively helical in the region spanning from G29 to G38 (all residues occupy the helical state to nearly 100%), the helix of A*β*_43_ is very distorted. The green plot on Figure 4a shows that even the more stable section between residues 30 and 36 is only 60-80% helical. The perturbation of the helicity is also indicated by the mean backbone (BB) hydrogen bond lengths (from the n*th* to the n-4*th* residue) plotted on Figure 4b.

If the information on figures 3 and 4 is combined, it is evident that the extend of sliding and unwinding is not the same in all processing steps: Cleaving C99 to A*β*_49_ causes strain in the N-terminal region (N27, K28 are less often helical) and unwinding at the C-terminus from residue 42 on. K28 on the other hand, edges just 1.8Å closer to the active site of PS, while the overall helicity decreases by 4.5% (see also table 3).

**Table 3:**
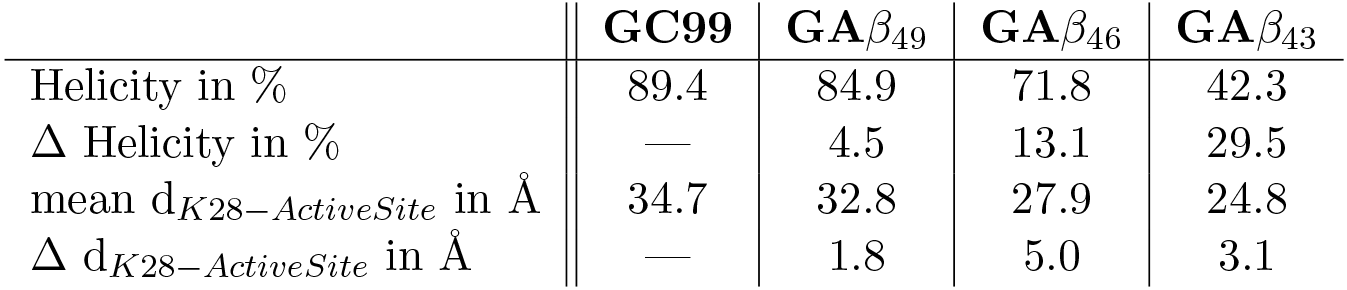
Mean helicity in % (calculated for residues D23 to T43 in all cases) and K28 – active site distances (in Å). The Δ – values denote changes from the n*th* to the n-1*st* cleavage step

The processing step leading to A*β*_40_, however, is above all characterized by a 5.0Å K28 displacement towards the PS TMDs. The mean helicity decreases by 13.2%, which is largely due to the helix opening in the C-terminus. The N-terminus is even slightly more helical than in simulation GA*β*_49_ (see Figure 4a).

The substrate helix is 29.2% less helical in GA*β*_43_ than it is in GA*β*_46_, while K28 moves 3.1Å closer to the active site in this processing step. Since the energetic cost to disrupt all the hydrogen bonds in an *α*-helix is significant (even though the open form is stabilized by water molecules present in the cavity), it begs the question why GA*β*_43_ does not just slide further towards the active site.

By analyzing the data gathered from the simulations, we identified two possible explanations:

The first one is the already discussed N-terminal anchor, which on one hand prevents complete dissociation of the E-S complex but on the other hand also keeps the N-terminus from sliding deeper into the membrane. Another explanation can be found by examining how A*β_n_* residues K28 and N27 interact with their environment:

In all conducted simulations, these residues, even though being close to GSEC, predominately interacted with the phosphate headgroups of POPC via Coulombic forces. This finding confirms the proposed role of K28 as an anchor in the lipid bilayer.^44^ According to our measurements, K28 is pulled inside the membrane helix while maintaining interactions with water molecules and the POPC headgroups. Due to the length of the side chain this seemed to work well for GA*β*_49_ and GA*β*_46_ but in GA*β*_43_, the amino group moved already towards the hydrophobic region, still maintaining polar interactions. This relocation distorted the membrane surface and pulled polar residues and water to the bilayer, thereby forming a hydrophilic sphere around the amino group of K28 (see figure 5, left panel). Evidently, this is energetically unfavorable and indicates that cleaving steps requiring such rearrangements are impaired by high reaction barriers.

**Figure 5:**
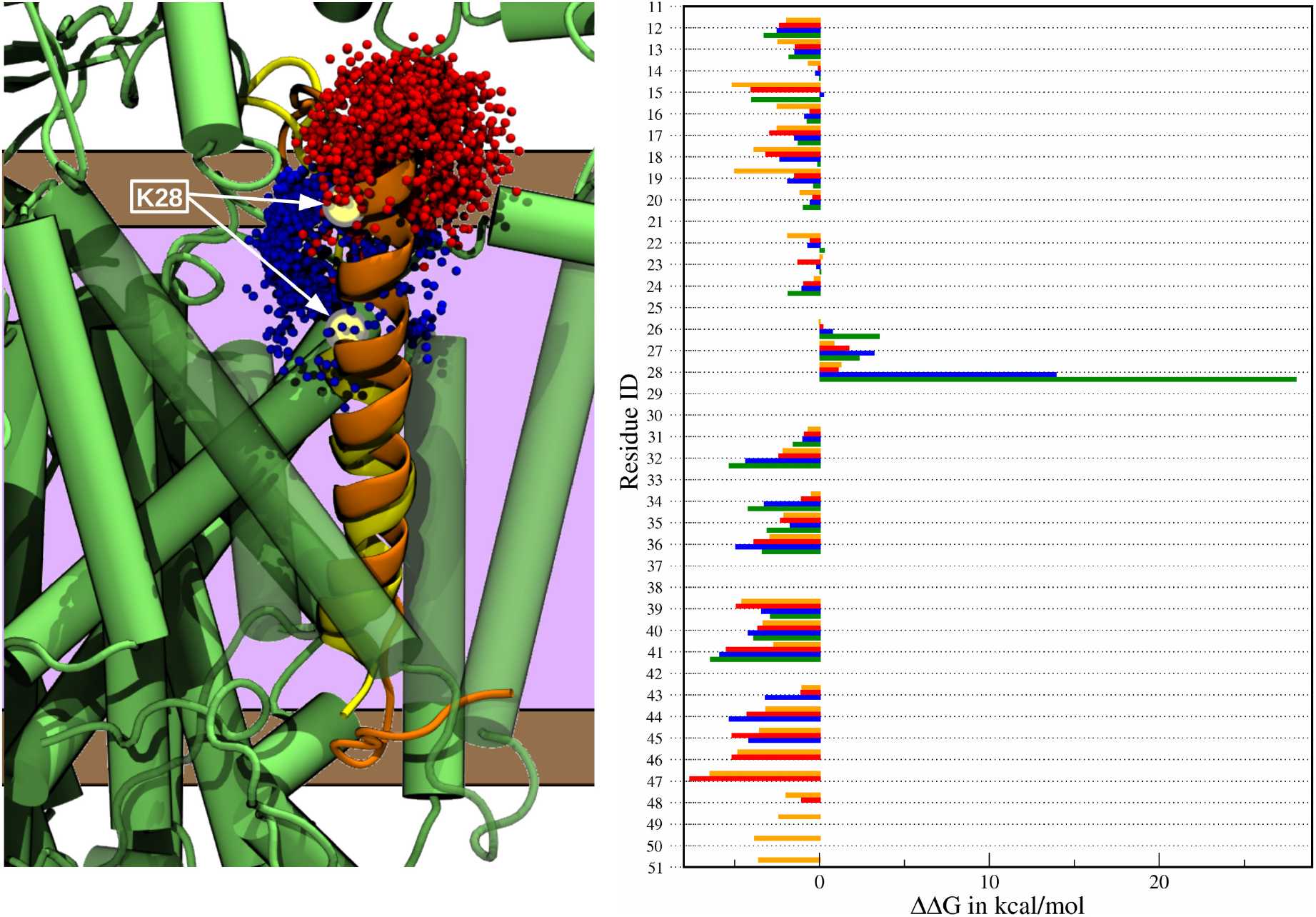
Left panel: Comparison of C99 (orange) and A*β*_43_ (yellow) in complex with GSEC (green). The membrane region is indicated in brown (headgroups) and purple (lipid tails). Water occupancy (superpositon of every tenth evaluation frame) close to K28 on C99 is shown as red spheres, while waters close to K28 on A*β*_43_ are colored blue. K28 has penetrated the lipid region in simulation GA*β*_43_, creating a hydrophilic pore. Right panel: In-silico alanine scans of all eligible residues from V12 to M51. Results for the GC99 case are shown as orange bars, GA*β*_49_ red, GA*β*_46_ blue and GA*β*_43_ green. A negative ΔΔG value indicates complex destabilization upon mutation into alanine, while a positive value signifies that the Ala-mutant interacts more favorably with the enzyme. The gaps in the data set are due to the fact that residues of type Gly or Ala can for methodological reasons, not be used for in-silico alanine scans.

The scissile bond has been pulled towards the active site via COM distance restraints in the simulations, therefore some interactions necessarily need to disrupt in order to allow close contacts at the active site. Our evaluation of the GA*β*_43_ simulation implies that the hydrogen bonding network in the helical region was weaker than the anchoring effect of K28 and instead of moving further into the cavity, the secondary structure of GA*β*_43_ began to change. Since this too involves a significant energetic penalty it seems plausible to conclude that it is a major reason why APP processing usually stops after cleaving to A*β*_40_.

To quantify the role of the juxtamembrane region of A*β_n_* peptides, we conducted MMPBSA based in-silico alanine-scans for all (eligible) amino acids present in our substrate model (Figure 5, right panel). Indeed, most substitutions to alanine are predicted to destabilize the complex. However, the polar and charged residues situated at the substrate’s juxtamemrane region clearly have an adverse effect on the stability of the GSEC-A*β_n_* complexes:

While K28 and N27 also destabilize the E-S complex in GC99 and GA*β*_49_, results for GA*β*_43_ and GA*β*_46_ predict a substantial increase in substrate-enzyme affinity for K28A and N27A mutations, suggesting that the insertion of the substrate molecule further into the membrane is energetically unfavorable. K28 acting as an anchor, necessitating higher activation energies and the fact that K28A mutations stabilize the complex (increasing E-S complex life time) may therfore explain why the K28A mutation in experimental studies leads to very short (<40AA) A*β* peptides.^44^

We were also able to investigate how complex formation impacts substrate helicity: According to our evaluations, shown on Figure 4a, free C99 (colored in black) remains helical up until residue L52. (while the secondary structure of K53 is more ambiguous). Bound to GSEC, however, C99 begins to unwind at T48 and by taking the standard deviations in Figure 4b into account as well, one can argue that helix destabilization begins already at I45.

This data is in good agreement with a recent NMR study of the transmembrane domain of APP^45^ that investigated the amide chemical shift perturbation caused by binding to the enzyme. The authors observed very subtle hydrogen bond destabilization beginning at V39, an increase at around V44 to T48 and a jump in chemical shift perturbation at residue L49. Qualitatively, this fits very well to the observed weakening of hydrogen bonds in our simulations. At the N-terminus, the study reported that A30 exhibited weaker hydrogen bonds in free C99 than in bound form. Figure 4b shows that this is also the case in our simulations. A similar finding, concerning residues G33 and L34 could not be replicated, since the hydrogen bonds were stable in both (GC99 and free C99) simulations.

The continuous unwinding of the substrate helix may be attributed to the presence of water molecules at the A*β_n_* C-terminal region. In order to test this hypothesis we have calculated the mean number of water molecules close to the backbone of residues V44 to L52 in simulations C99 and GC99 or the nine C-terminal residues in simulations GA*β*_49_, GA*β*_46_ and GA*β*_43_, respectively (Table 4). The results indicate that the number of water molecules is lower in the simulation of free C99, compared to the complex bound forms. This is due the fact that the binding site of C99 which is situated within the TMDs of PS acts like a pore that allows water molecules to enter the otherwise hydrophobic intra-membrane region. The hydration numbers suggest that helix unwinding is at least partially mediated by the presence of water molecules in the active site of GSEC. Binding to the enzyme, however, can also be expected to have a distinct effect on the helicity of the substrate.

**Table 4:** Mean number of water molecules within 5Å of the C-terminal A*β_n_* backbone in the respective simulations.

| Simulation | Residues | #Water | Std. Dev. | Min. | Max. |
| --- | --- | --- | --- | --- | --- |
| free C99 | V44 to L52 | 9.0 | 3.2 | 0 | 21 |
| GC99 | V44 to L52 | 14.0 | 2.8 | 6 | 24 |
| GA$\beta_{49}$ | I41 to L49 | 14.2 | 2.3 | 7 | 22 |
| GA$\beta_{46}$ | G38 to V46 | 17.5 | 3.7 | 8 | 29 |
| GA$\beta_{43}$ | M35 to T43 | 17.2 | 2.6 | 11 | 28 |

### 1.4 A*β_n_* is in close contact with many known FAD-mutation sites

As already established, A*β_n_* is in proximity to a large number of FAD-mutation sites located on PS when binding to the cavity formed by TMDs 2 3 and 5. For further investigation, we re-evaluated the mutation site proximities for the more representative simulations GC99, GA*β*_49_, GA*β*_46_ and GA*β*_43_ where the complexes are in transition state – like conformations with active– and cleavage-sites in contact. Here, we found even more mutation sites to be within 5Å of the substrate helices (mean values over 1500 frames): 38 in GC99, 28 in GA*β*_49_, 24 in GA*β*_46_ and 23 in GA*β*_43_. An overview over the nature and position of the mutation sites is given in table 5, from where also the abundance of proximity can be taken (given in percent of evaluation frames). A depiction of C99 in complex with GSEC and surrounded by FAD – mutation sites can be found in the supplement.

**Table 5:**
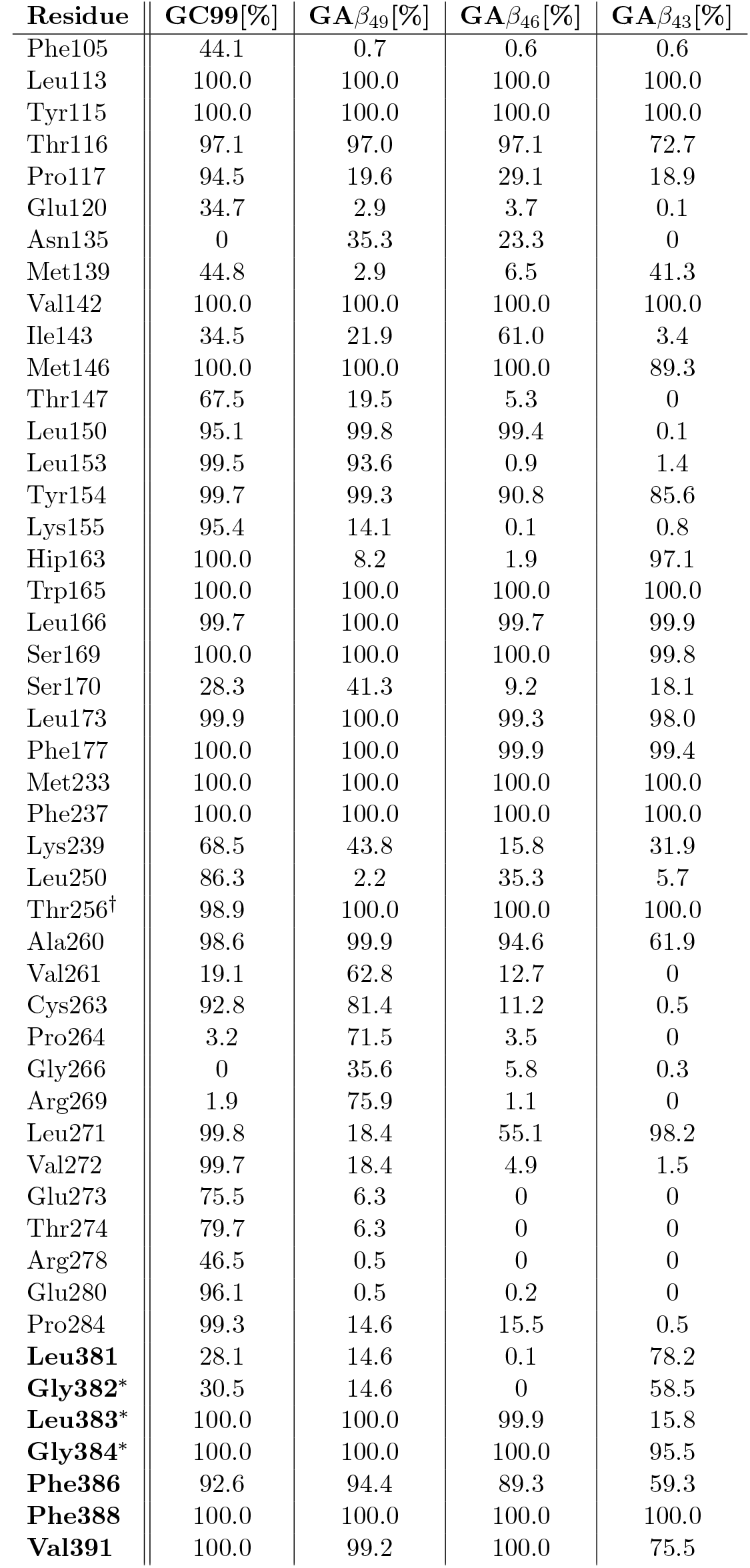
Known FAD-causing mutation sites that are in close contact with A*β_n_*. Only mutation sites that are in at least 30% of evaluation frames within 5Å of a substrate residue in at least one of the simulations are shown. The given percentages correspond to the number of evaluation frames in which the respective residue was within 5Å of the substrate. * Part of the GXGD motif (L383 is not a known FAD mutation site, it has been shown, however, that L383X mutations influence GSEC behavior as well^26^). † The 5FN2 PDB stucture that was used as starting template for the simulations featured a Y256T mutation. Residue names printed in bold font are situated on the C-terminal fragment of PS.

| Residue | GC99[%] | GA$\beta_{49}$[%] | GA$\beta_{46}$[%] | GA$\beta_{43}$[%] |
| --- | --- | --- | --- | --- |
| Phe105 | 44.1 | 0.7 | 0.6 | 0.6 |
| Leu113 | 100.0 | 100.0 | 100.0 | 100.0 |
| Tyr115 | 100.0 | 100.0 | 100.0 | 100.0 |
| Thr116 | 97.1 | 97.0 | 97.1 | 72.7 |
| Pro117 | 94.5 | 19.6 | 29.1 | 18.9 |
| Glu120 | 34.7 | 2.9 | 3.7 | 0.1 |
| Asn135 | 0 | 35.3 | 23.3 | 0 |
| Met139 | 44.8 | 2.9 | 6.5 | 41.3 |
| Val142 | 100.0 | 100.0 | 100.0 | 100.0 |
| Ile143 | 34.5 | 21.9 | 61.0 | 3.4 |
| Met146 | 100.0 | 100.0 | 100.0 | 89.3 |
| Thr147 | 67.5 | 19.5 | 5.3 | 0 |
| Leu150 | 95.1 | 99.8 | 99.4 | 0.1 |
| Leu153 | 99.5 | 93.6 | 0.9 | 1.4 |
| Tyr154 | 99.7 | 99.3 | 90.8 | 85.6 |
| Lys155 | 95.4 | 14.1 | 0.1 | 0.8 |
| Hip163 | 100.0 | 8.2 | 1.9 | 97.1 |
| Trp165 | 100.0 | 100.0 | 100.0 | 100.0 |
| Leu166 | 99.7 | 100.0 | 99.7 | 99.9 |
| Ser169 | 100.0 | 100.0 | 100.0 | 99.8 |
| Ser170 | 28.3 | 41.3 | 9.2 | 18.1 |
| Leu173 | 99.9 | 100.0 | 99.3 | 98.0 |
| Phe177 | 100.0 | 100.0 | 99.9 | 99.4 |
| Met233 | 100.0 | 100.0 | 100.0 | 100.0 |
| Phe237 | 100.0 | 100.0 | 100.0 | 100.0 |
| Lys239 | 68.5 | 43.8 | 15.8 | 31.9 |
| Leu250 | 86.3 | 2.2 | 35.3 | 5.7 |
| Thr256<sup>†</sup> | 98.9 | 100.0 | 100.0 | 100.0 |
| Ala260 | 98.6 | 99.9 | 94.6 | 61.9 |
| Val261 | 19.1 | 62.8 | 12.7 | 0 |
| Cys263 | 92.8 | 81.4 | 11.2 | 0.5 |
| Pro264 | 3.2 | 71.5 | 3.5 | 0 |
| Gly266 | 0 | 35.6 | 5.8 | 0.3 |
| Arg269 | 1.9 | 75.9 | 1.1 | 0 |
| Leu271 | 99.8 | 18.4 | 55.1 | 98.2 |
| Val272 | 99.7 | 18.4 | 4.9 | 1.5 |
| Glu273 | 75.5 | 6.3 | 0 | 0 |
| Thr274 | 79.7 | 6.3 | 0 | 0 |
| Arg278 | 46.5 | 0.5 | 0 | 0 |
| Glu280 | 96.1 | 0.5 | 0.2 | 0 |
| Pro284 | 99.3 | 14.6 | 15.5 | 0.5 |
| **Leu381** | 28.1 | 14.6 | 0.1 | 78.2 |
| **Gly382\*** | 30.5 | 14.6 | 0 | 58.5 |
| **Leu383\*** | 100.0 | 100.0 | 99.9 | 15.8 |
| **Gly384\*** | 100.0 | 100.0 | 100.0 | 95.5 |
| **Phe386** | 92.6 | 94.4 | 89.3 | 59.3 |
| **Phe388** | 100.0 | 100.0 | 100.0 | 100.0 |
| **Val391** | 100.0 | 99.2 | 100.0 | 75.5 |

The simulations suggest that in our model a very large number of crucial PS residues is close to the substrate during all steps of proteolytic processing. Naturally, C99 being longer than the processing intermediates is in contact with more mutation sites than the latter. However, the large number of shared sites indicates that the processing pathway may not exclusively be determined at the endo-peptide stage but also during *γ* – or *ζ*-cleavage.^19^ In addition to FAD-causing mutation sites, also the GXGD motif is in proximity of a number of A*β_n_* residues. Extensive mutagenesis studies^25^’^26^’^46^ have shown that increasing the size of the Gly residues and/or mutation of the x-residue in the GXGD motif has a severe impact on GSEC processivity, pathogenicity a well as substrate selectivity. This hints at the existence of key interactions between enzyme and substrate in this area.

Recently, Fukumori et al.^28^ performed photoaffinity mapping experiments and reported interactions between nearly all C99 residues and the N-terminal fragment of PS (TMDs 1 to 6). This fits very well to our proposed binding mode, where also the N-terminal fragment of PS presents the majority of interfacing residues. Additionally, Fukumori et al. were able to cross link C99 to the C-terminal PS fragment. Especially C99 residues 51, 52 and 54 have been found to be major cross-linking sites when the natural amino acids were replaced by 4-Benzoyl-L-phenylalanine (Bpa). In our simulations these residues are close enough to bind to the C-terminal fragment. Based on our model, the most likely interaction sites on the C-terminal fragment for these residues are: P432 (for M51), K380 (L52), D385 (L52) and L383 (K54). The majority of sampled frames shows these amino acids to be in contact with the N-terminus, however, Fukumori et al. have found that these residues can be cross-linked to the N-terminal fragment as well.

Besides PS, also interactions with NCT and PEN-2 have been reported. We find our study to be in excellent agreement also regarding NCT interactions for all the residues that cross link and have been included in our model (residues F19 and A21). Both are constantly in contact with NCT, where F19 binds to I246 and R652, while A21 is very close to residues N243, P244 and E245. No interactions with Pen-2 could be identified in our simulation, indicating that the associated binding events that have been reported by Fukumori et al. are of significance for substrate recruitment rather than the actual processing step.

The GC99 E-S complex model is also able to explain the findings of photolabeling and binding competition experiments performed by Kornilova et al.:^47^ While a 10 AA long helix derived from the C99 sequence (D-10) which is a known inhibitor of GSEC has been shown bind to the enzyme even in the presence of transition state analogue inhibitors, a 13 AA long helix (D-13, also an inhibitor) competes with active site directed compounds. The authors offered two possible explanations for their findings: “(i) D-13 binds to the initial binding site but protrudes into the active site because of its three additional amino acids; or (ii) it binds to the initial binding site and allosterically affects the active site by means of the interaction of its three extra amino acids with the protein.”^47^

Figure 6, a snapshot from the GC99 simulation, indicates the putative positions of the D-10 and D-13 helices based on the locations of the corresponding amino acids in C99. While the C-terminus of the shorter D-10 helix (blue in fig. 6) is still relatively far away from the active site, allowing for enough space for an active site directed compound to bind, D-13 (magenta) is already in close contact with D385 and is expected to compete with binding of transition state analogues to the active site. Hence, the predicted binding model supports the former hypothesis of Kornilova et al.^47^ that D13 steriochemically interferes with transition state inhibitor binding and offers an explanation based on the positioning of C99.

**Figure 6:**
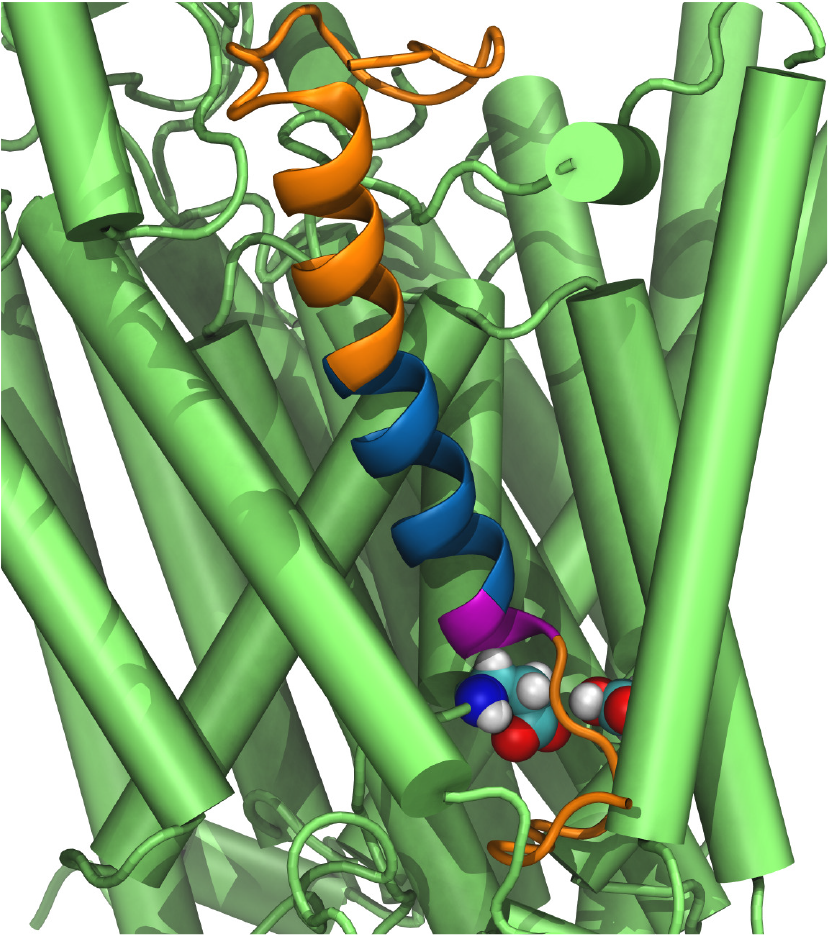
Illustration of the putative positions of designed D-10 (blue) and D-13 (additional three amino acids in magenta) helices by Kornilova et al.^47^ aligned to the GC99 model (C99 in orange). The PS-1 active site aspartates are shown as Van-der-Waals spheres.

Due to rotation symmetry, the orientation of the substrate relative to the enzyme is a crucial aspect of any binding model involving *α*-helices. The detailed study on APP mutations performed by Xu et al.^18^ allowed us to test the validity of the C99-GSEC interface found in our model. Among other aspects, Xu et al. assessed the impact of known C99-FAD mutations on the efficiency of *ϵ*-cleavage. It was found that certain mutations of D7, E11, K16, E22, D23, L34, A42, T43, V44, I45, V46, T48, L52 and K53 severely reduced the activity of *ϵ*-cleavage. These results are in very good agreement with our binding model, since all of the applicable (D7 and E11 are not part of our C99 model) mutation sites are in contact with either nicastrin or presenilin (see supplement for details on interactions). The only exception is L34 which faces away from GSEC in our binding model. However, Leu to Val mutations have a known adverse effect on helix formation tendency^48–50^ and it can be expected that interactions of both amino acids with PS would be very similar. Therefore it is quite likely that the measured impact on *ϵ*-cleavage proficiency stems from changes of helicity of the substrate, rather than interactions with the enzyme, hence no direct contact between L34 and GSEC is required to explain the effect of this mutation. Additionally, Xu et al. investigated also non (or at least of unknown effect)-FAD mutations and found that especially mutations of T43, V46, T48 and L49 to very bulky amino acids (Trp, Tyr or Arg) have a very detrimental influence to the efficiency of *ϵ*-cleavage. In the presented binding model all of these residues are in very close contact with the enzyme, offering an explanation for each mutation: V46 and L49 are in close proximity to the crucial GXGD motif and T43 binds in a pocket formed by three known FAD mutation sites on PS: M233 (TMD 5), F237 (TMD 5) and V391 (TMD 7). Thr48, on the other hand is buried in a very stable pocket and is also in contact with PS-FAD mutation sites (V142 and M146).

Overall, the presented model features many close substrate-enzyme interactions at sites that are known to be crucial for the cleaving of C99, lending credibility to its ability to elucidate how PS mutations impact the catalytic properties of GSEC.

## 2 Conclusions

We have performed twelve *μ*s-timescale simulations of free C99, as well as various GSEC-A*β_n_* complexes and identified the binding cavity formed by PS TMDs 2, 3 and 5 to be the most likely binding site for the substrate. In this pocket, the substrate is in contact with the N– and C– terminal fragments of PS and additionally, with the extracellular region of nicastrin. Furthermore, the simulations and subsequently performed MMPBSA free energy estimations suggested that NCT acts as a binding partner for A*β_n_* during processing.

In addition to the C99-GSEC complex, also the processing intermediates A*β*_49_, A*β*_46_ and A*β*_43_ have been investigated. This allowed us to draw conclusions about the possible mechanism of substrate repositioning in the active site and it was revealed that a combined unwinding/sliding mechanism is responsible for repositioning the scissile bond to induce subsequent hydrolysis steps. Regarding the mechanism, we also found that even though large parts of the helices are repositioned/unwound during processing, many residues belonging to the N-terminal loop region of the A*β_n_* substrate peptides maintained their interactions with PS/GSEC throughout all simulations. This finding strongly supports the existence of a permanent N-terminal anchoring region as has been suggested previously.^13^ At least from a mechanistic point of view this binding area is an interesting candidate for modulator binding, since it remains topologically unaltered for the duration of C99 processing.

Based on previously existing knowledge and findings in this study, we propose that there are four different kinds of GSEC/APP mutations modulating the outcome of C99 processing: Mutations that (de)stabilize the GSEC-A*β_n_* complex, situated at the interface between enzyme and substrate. These mutations influence E-S complex lifetimes which in turn leads to Aβ peptides of different length. (2) Mutations on either GSEC or APP, leading to a shift towards a production line terminating at A*β*_42_ which dissociates from the enzyme before it can be cleaved to A*β*_39_. This work indicates that the dissociation is likely caused by reduced N-terminal interactions, strain of the remaining helix and the energetically unfavorable pulling of polar species inside the membrane region. (3) Mutations of K28/N27 on C99 to apolar species have been shown to lead to exceptionally short products.^44^ Our investigations suggest (and confirm) that K28/N27 anchor A*β_n_* helices to the headgroup region of the bilayer (a depiction of K28 interacting with two POPC molecules is provided in the supplement). During processing these very polar/charged side chains are pulled inside the membrane region which is energetically unfavorable. Removing these barriers aids the repositioning of very short intermediates and thereby increases the reaction rates of the later stages of C99 processing. Mutations on PS that lead to a change of processivity by impacting the probability of PS to occupy the catalytically active conformation, hence changing the intervals between two cleavage steps.

Another topic controversially discussed in the context of APP processing is APP homodimerization.^51–53^ Our study strongly disfavors the proteolytic cleavage of APP homodimers, as the binding model suggests that there is not enough space at the active site cavity to accommodate such dimers.

In this study we propose a working model of the APP-GSEC complex in atomistic detail. The presented binding mode is based mainly on three known structural properties: The apo-structure of *γ*-secretase,^4^ an NMR structure of C99^37^ and that at some point of substrate binding the scissile bond has to be in contact with the active site. Although other putative complex geometries cannot be completely excluded, the present working model is able to confirm and explain several well established experimental findings. Hence, the model can serve to guide mutagenesis, disulfide– and other cross-linking studies of the A*β_n_*-GSEC complex and encourage investigators to use the model to challenge and improve the proposed binding mode of the *γ*-secretase– A*β_n_* complex.

## 3 MATERIALS AND METHODS

In the course of this study, 12 simulations of C99 and C99-, A*β*_49_-, A*β*_46_-, A*β*_43_-GSEC complexes have been performed. An overview is provided in Table 6.

**Table 6:**
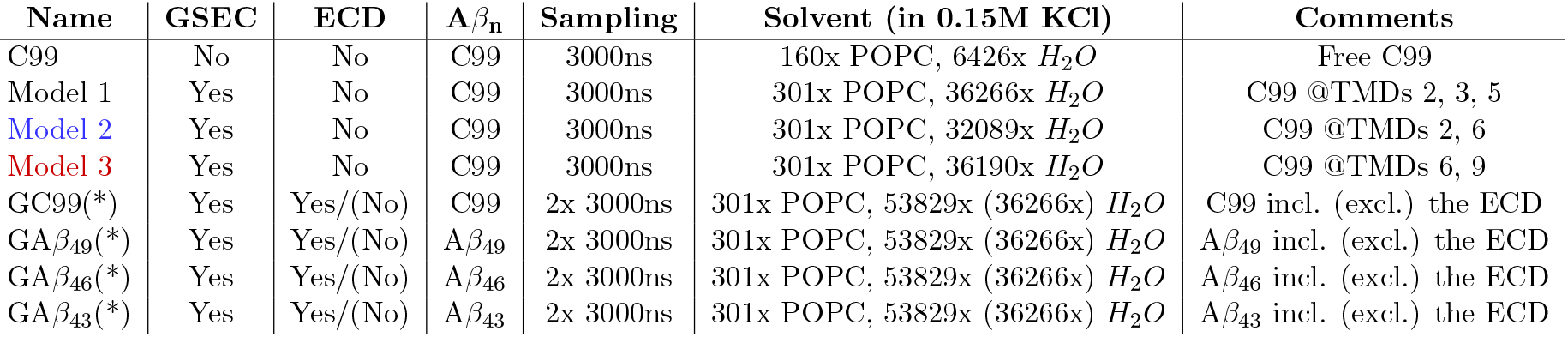
Overview over the simulations that have been conducted in the course of this investigation. “GSEC” indicates if the TMDs of GSEC were present in the simulation box, while ECD indicates the presence (or absence) of the NCT ECD. The parenthesis indicate that the simulation has been carried out twice, once in presence of the ECD and a second time in its absence (the absence of the ECD is denoted by a ‘*’ in the simulation name).

| Name | GSEC | ECD | A$\beta_n$ | Sampling | Solvent (in 0.15M KCl) | Comments |
| --- | --- | --- | --- | --- | --- | --- |
| C99 | No | No | C99 | 3000ns | 160x POPC, 6426x $H_2O$ | Free C99 |
| Model 1 | Yes | No | C99 | 3000ns | 301x POPC, 36266x $H_2O$ | C99 @TMDs 2, 3, 5 |
| Model 2 | Yes | No | C99 | 3000ns | 301x POPC, 32089x $H_2O$ | C99 @TMDs 2, 6 |
| Model 3 | Yes | No | C99 | 3000ns | 301x POPC, 36190x $H_2O$ | C99 @TMDs 6, 9 |
| GC99(*) | Yes | Yes/(No) | C99 | 2x 3000ns | 301x POPC, 53829x (36266x) $H_2O$ | C99 incl. (excl.) the ECD |
| GA$\beta_{49}$(*) | Yes | Yes/(No) | A$\beta_{49}$ | 2x 3000ns | 301x POPC, 53829x (36266x) $H_2O$ | A$\beta_{49}$ incl. (excl.) the ECD |
| GA$\beta_{46}$(*) | Yes | Yes/(No) | A$\beta_{46}$ | 2x 3000ns | 301x POPC, 53829x (36266x) $H_2O$ | A$\beta_{46}$ incl. (excl.) the ECD |
| GA$\beta_{43}$(*) | Yes | Yes/(No) | A$\beta_{43}$ | 2x 3000ns | 301x POPC, 53829x (36266x) $H_2O$ | A$\beta_{43}$ incl. (excl.) the ECD |

### 3.1 Starting structures

The *γ*-secretase structures used in the simulations have been taken from already equilibrated structures of the *μ*s – timescale production trajectories of a study on the apo-structure of GSEC,^30^ which in turn were based on the PDB structure 5FN2^4^ (RMSDs of the apo-state simulations can be found in the supplement). The starting conformation of the C99 fragment (residues V12 to Y57) used in this investigation has been taken from PDB file 2LP1.^37^ The substrate fragment used in the simulations is considerably longer than one of the shortest C99 chains that are known to be processed by GSEC (E22 to K55).^54^

The simulation box of the C99 simulation has been created by first submitting the PDB file to the PPM-^55^ and subsequently to the CHARMM-GUI – server^56,57^ to insert the lipid spanning part of C99 in a physically reasonable way into a POPC membrane. The protonation states of the titratable side chains were chosen according to their natural state at pH 6.5.

In a pre-equilibration step, target temperature and pressure were reached in a step-wise pre-equilibration phase, as suggested by the CHARMM-GUI server (details can be found in Supporting Information). After 500ns of equilibration, a 3000ns sampling trajectory was generated.

ACE and NME capping groups have been used for the N– and C-terminus of C99 in simulations C99, Model 1, Model 2, Model 3, GC99 and GC99*. In all other simulations, only the N-term of the A*β_n_* chain has been capped. In simulations containing GSEC, the N-termini of NCT, PS-1 and APH-1 were capped with ACE groups, while the C-termini of NCT and APH-1a were terminated via NME. In all other cases the number of missing residues was so low that a charged tail was deemed to be more realistic than a capped one.

### 3.2 Generation of the GSEC-A*β_n_* complexes

The Model 1, 2 and 3 structures were generated by placing the equilibrated C99 chain close to the investigated binding surface on the outside of PS. The GSEC structure was taken from an equilibrated (3500ns long) simulation of the apo-conformation of GSEC, missing the ECD.^30^

Aiier a 100ns long equiliuraiion phase (lonow-ing the same pre-equilibration procedure as described above), the *ϵ*-site on C99 (residues 49 and 50) was pulled towards the active site of PS in 1 A steps within a 100ns time-frame using harmonic center of mass (COM) restraints (*k* = 5 *kcal* * *mol^−1^* *Å^−2^) in order to bring the active site of PS in close contact with the substrate’s scissile bond. Subsequently, the restraints were removed and the simulation was allowed to equilibrate freely after which a 3000ns long evaluation trajectory was sampled.

The starting structure for simulation GC99* (the ‘*’ denotes the missing ECD) of the GSEC-C99 complex was prepared by decreasing the distance between TMDs 2 and 3 with the use of weak COM restraints (*k* = 0.5 *kcal* * *mol^−1^* *Å^−2^) for 100ns. This was necessary because the insertion of C99 caused the increase of the TMD 2-TMD distance to allow passage of the substrate into the binding site. This circumstance also confirms the putative role of TMD 2 as a flexible entry gate for the substrate helices.

After 1000ns of free simulation, since the Model 1, 2, 3 simulations showed that the *ϵ*-site does not stay in contact with the active site for a substantial amount of time, we forced the groups that would be involved in the hydrolysis into a transition-state like conformation^58^

(see also figure 7). This enabled us to study the GSEC-C99 complex in the state in which it could also be cleaved. Therefore, the active site restraints remained active for the complete sampling phase (which again followed 500ns of equilibration).

**Figure 7:**
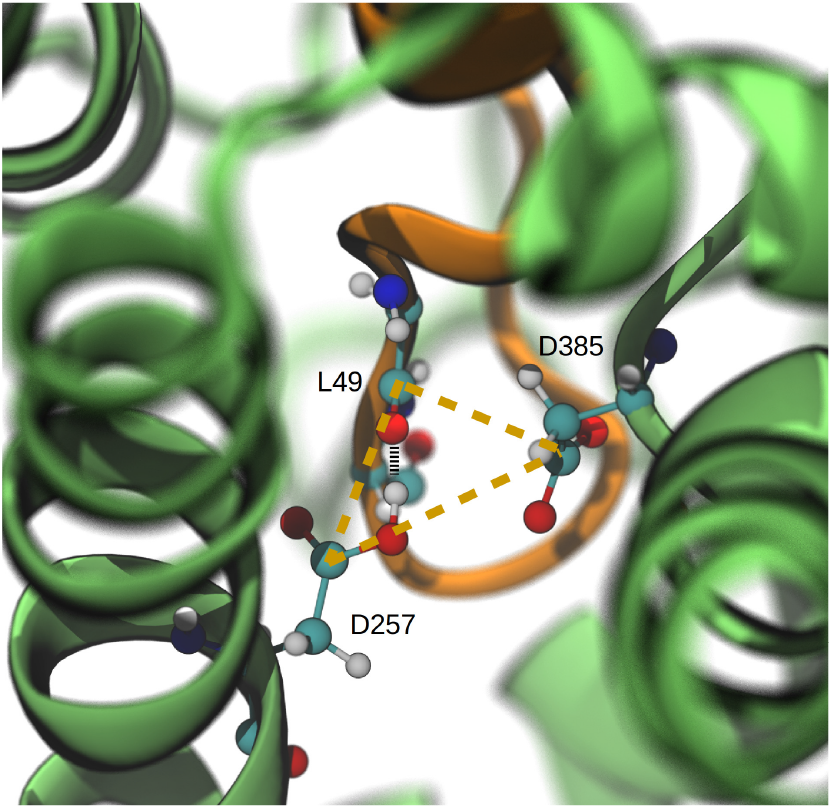
Transition state-like conformation at the active site that was enforced in simulations GC99(*), GA*β*_49_(*), GA*β*_46_(*) and GA*β*_43_(*). Residue D257 (left) is situated at TMD 6, while D385 (right) is part of TMD 7. The arrangement was stabilized by appropriate distance restraints between carbonyl group of the scissile bond and atoms of residue D257 and D385.

The starting structure of the GC99 simulation (including the ECD) was constructed by taking the PS-C99 complex (discarding the rest of GSEC) from the Model 1 complex simulation and merging it with full GSEC. The full GSEC structure was taken from a snapshot of a 1000ns long simulation of the complete GSEC complex.^30^ The two structures were merged by replacing the original apo-PS with the PS-C99 complex from the Model 1 simulation. Starting from this merged structure, the simulation protocol was exactly the same as for GC99*. In both types of simulations – with and without the ECD, the shortened A*β_n_* chains were generated by deleting the residues that are removed by the enzymatic hydrolysis. Again, the systems (GA*β*_49_, GA*β*_4_6, GA*β*_43_) were allowed to equilibrate for 1000ns and subsequently forced into transition-like states. These states were simulated for 3500ns and after discarding the first 500ns, the trajectories were used for evaluation. All proteins have been simulated in a 0.15M KCl solution to ensure a realistic, physiological environment.

### 3.3 Simulation Details

The simulations have been performed using the CUDA^59^ accelerated version of PMEMD^60–62^ part of the AMBER16 package.^63^ Proteins and lipids have been described by the AMBER14SB^64^ and Lipid14^65^ force fields respectively, while the TIP3P^66^ model has been used for water. The cut-off for Coulombic and other non-bonded forces were set to 8 A, all interactions beyond that were described by the particle mesh ewald method.^67^ Periodic boundary conditions were used, with box sizes ranging from ~ 400000 (C99) over ~ 1700000 (Model 1, 2, 3) to ~ 2200000 Å^3^ (GC99, GA*β*_49_, GA*β*_46_, GA*β*_43_). The target temperature for all simulations was 303.15K, using the Langevin thermostat^68^ with a collision frequency of 1ps^−1^. The Monte Carlo barostat^69^ was used to generate NPT ensembles with a target pressure of 1.0 bar. In order to achieve time steps of 4.0fs, the SHAKE^70^ algorithm has been utilized in conjunction with hydrogen mass repartitioning.^71^ Trajectory analysis was carried out using CPPTRAJ^72^ (also part of the AMBER16 package) and VMD,^73^ which was also used for visualizing the output and rendering the snapshots. We included further details regarding the COM restraints and pre-equilibration protocol in the supplement. With average C*α* RMSDs (disregarding the highly mobile loop region between TMDs 6 and 7 in all simulations including GSEC) ranging from 2.4Å (GA*β*_43_*) to 4.1Å (GC99) all simulated structures remained stable throughout the sampling phase (RMSD plots for all simulations can be found in the supplement).

### 3.4 Free Energy Calculations

Free energy calculations and alanine-scans have been conducted using the single-trajectory MMPBSA^74^ post-processing method, utilizing the MMPBSA.py^75^ program.

An implicit membrane model has been employed in order to describe the system electrostatics as realistically as possible. The membrane has been assumed to be 34 Å thick. This made sure that the permittivity of the region that was originally spanned by the POPC lipid tails, could be set to *ε* = 2.0 in the Poisson-Boltzmann implicit solvation treatment.^76^

Our simulations showed that a large number of water molecules were level with the headgroup region of the membrane – especially in the cavity between PS and the GSEC ECD (which is void of lipids), therefore the phosphatidylcholine – region was included in the the aqueous layer (with *ε* = 80.0) of the model to ensure realistic electrostatic interactions between the N-terminus of A*β_n_* and GSEC. The internal permittivity (of the protein) has been assumed to be *ε* = 1.0.

The center of the membrane slab (in z-direction) was set to the mean value of the center of mass of the POPC bilayer in the respective trajectory and periodic boundary conditions have been used to ensure the a realistic treatment of the membrane. The ionic strength was set to be equal to the one in the explicit treatment (0.15M).

In every MMPBSA calculation, 1500 frames have been evaluated in order to generate reliable statistics. Due to the enormous system size, the computational expense of the implicit membrane treatment and the large number of MMPBSA calculations that were necessary for this study, no entropy corrections (beyond the intrinsic PBSA entropy terms) have been performed. Therefore, the presented energies cannot be directly compared to experimentally derived ΔG values.

## Supporting Information

Additional pictures and simulation details are supplied in pdf format.

## List of Abbreviations

AA – amino acid A*β* – Amyloid *β*

ACE – acetyl AICD – amyloid intracellular c-terminal domain

APH-1 – anterior-pharynx-defective-1

COM – center of mass

GSEC – *γ*-secretase

APP – amyloid precursor protein

ECD – ectodomain

E-S complex – enzyme – substrate complex

FAD – familial Alzheimer’s disease

MD – molecular dynamics

MMPBSA – molecular mechanics Poisson-Boltzmann surface area

NCT – nicastrin

NME – N-methyl Pen-2 – presenilin enhancer 2

PS – presenilin

RMSD – root mean square deviaton

SASA – solvent accessible surface area

TMD – trans membrane domain

## Author Informations

M.H. performed research, analyzed data, and wrote the article; and M.Z. designed research and wrote the article.

## Acknowledgments

We thank L. Chávez-Gutiérrez and H. Steiner for many helpful discussions. Financial support by the DFG (German Research Foundation) grant FOR 2290 (project P7) is gratefully acknowledged. Computer resources for this project have been provided by the Gauss Centre for Supercomputing/Leibniz Supercomputing Centre under grant pr27za.

## Notes

#### Summary of Updates

Figures 1 and 3 have been revised. Additionally, some stylistic problems and typos have been taken care of. An additional picture has been added to the supplement.

## References

[1] D. Langosch, C. Scharnagl, H. Steiner, and M. K. Lemberg. Understanding intramembrane proteolysis: from protein dynamics to reaction kinetics. Trends Biochem. Sci., 40:318–327, 2015.

[2] D. Langosch and H. Steiner. Substrate processing in intramembrane proteolysis by – secretasethe role of protein dynamics. Biol. Chem., 398:441–453, 2017.

[3] X. Bai, C. Yan, G. Yan, P. Lu, D. Ma, L. Sun, R. Zhou, S. H. W. Scheres, and Y. Shi. An atomic structure of human *γ*-secretase. Nature, 525:212–218, 2015.

[4] X. Bai, E. Rajendra, G. Yuang, Y. Shi, and S. H. W. Scheres. Sampling the conformational space of the catalytic subunit of human *γ*-secretase. eLife, 4:pii:e11182, 2015.

[5] M. S. Wolfe, W. Xia, B. L. Ostaszewski, T. S. Diehl, Taylor K. W, and D. J. Selkoe. Two transmembrane aspartates in presenilin-1 required for presenilin endoproteolysis and *γ*-secretase activity. Nature, 398:513, 1999.

[6] Y. Zhang, W. Luo, H. Wang, P. Lin, K. S. Vetrivel, F. Liao, F. Li, P. C. Wong, M. G. Farquhar, G. Thinakaran, and X. Huaxi. Nicastrin is critical for stability and trafficking but not association of other presenilin/*γ*-secretase components. J. Biol. Chem., 280:17020–17026, 2005.

[7] R. Francis, G. McGrath, J. Zhang, D. A. Ruddy, M. Sym, J. Apfeld, N. Javier, M. Monique, M. Maxwell, B. Hai, M. C. Ellis, A. L. Parks, W. Xu, J. Li, M. Gurney, R. L. Myers, C. S. Himes, R. Hiebsch, C. Ruble, J. S. Nye, and D. Curtis. aph-1 and pen-2 are required for notch pathway signaling, *γ*-secretase cleavage of βapp, and presenilin protein accumulation. Dev. Cell, 3:85–97, 2002.

[8] S. F. Lee, S. Shah, C. Yu, W. C. Wigley, H. Li, M. Lim, K. Pedersen, W. Han, P. Thomas, J. Lundkvist, Y-H. Hao, and G. Yu. A conserved gxxxg motif in aph-1 is critical for assembly and activity of the *γ*-secretase complex. Journal of Biological Chemistry, 279(6):4144–4152, 2004.

[9] R. Vassar, B. D. Bennett, S. Babu-Khan, S. Kahn, E. Mendiaz, P. Denis, D. B. Teplow, S. Ross, P. Amarante, and R. Loeloff et al. β-secretase cleavage of alzheimer’s amyloid precursor protein by the transmembrane aspartic protease bace. Science, 286:735–741, 1999.

[10] H. Steiner, A. Fukumori, S. Tagami, and M. Okochi. Making the final cut: pathogenic amyloid-peptide generation by -secretase. Cell Stress, 2:292–310, 2018.

[11] Y. Zhang, R. Thompson, H. Zhang, and H. Xu. App processing in alzheimer’s disease. Mol. Brain, 4:3, 2011.

[12] O. Holmes, S. Paturi, Y. Wenjuan, M. S. Wolfe, and D. J. Selkoe. Effects of membrane lipids on the activitiy and processivity of purified *γ*-secretase. J. Biochem., 52:3565–3575, 2012.

[13] M. Szaruga, B. Munteanu, S. Lismont, S. Veugelen, K. Horre, M. Mercken, T. C. Saido, N. S. Ryan, T. De Vos, S. N. Savvides, R. Gallardo, J. Schmykowitz, F. Rousseau, N. C. Fox, B. De Strooper, and L. Chávez-Gutiérrez. Alzheimer’s-causing mutations shift a*β* length by destabilizing *γ*-secretase A*β*n interactions. Cell, 170:443–456, 2017.

[14] J. Hardy and D. Allsop. Amyloid depoistion as the central event in the aetiology of alzheimer’s disease. Trends Pharamcol., 12:383–388, 1991.

[15] T. J. Jarrett, E. P. Berger, and P. T. Lansbury Jr. The c-terminus of the *β* protein is critical in amyloidogenesis. Ann. N.Y. Acad. Sci., 695:144–148, 1993.

[16] T. Iwatsubo, A. Okada, N. Suzuki, H. Mizusawa, N. Nukina, and Y. Ihara. Visualization of a*β*42 (43) and a*β*40 in senile plaques with end-specific a*β* monoclonals: evidence that an initially deposited species is a*β*42 (43). Neuron, 13:45–53, 1994.

[17] A. Burns and S. Iliffe. Alzheimer’s disease. Br. Me. J., 338:b158, 2009.

[18] T. Xu, Y. Yan, Y. Kang, Y. Jiang, K. Melcher, and H. E. Xu. Alzheimer’s disease-associated mutations increase amyloid precursor protein resistance to *γ*-secretase cleavage and the a*β*_40_/a*β*_42_ ratio. Cell Discovery, 1:16026, 2016.

[19] D. M. Bolduc, D. R. Montagna, M. C. Seghers, M. S. Wolfe, and D. J. Selkoe. The amyloid beta forming tripeptide cleavage mechanism of *γ*-secretase. eLife, 5:e17578, 2016.

[20] L. Chávez-Gutiérrez, L. Been, I. Benilova, A. Vandersteen, M. Benurwar, M. Borgers, S. Lismont, L. Zhou, S. Van Cleynenbreugel, H. Esselmann, J. Wiltfang, L. Serneels, E. Karran, H. Gijsen, J. Schymkowitz, F. Rousseau, K. Broersen, and B. De Strooper. The mechanism of *γ*-secretase dysfunction in familial alzheimer disease. EMBO J., 31:2261–2274, OPTurl =, OPTlastchecked =, OPTdoi =, 2012.

[21] Y. Qi-Takahara, M. Morishima-Kawashima, Y. Tanimura, G. Dolios, N. Hirotani, Y. Horikoshi, F. Kametani, M. Maeda, T. C. Saido, R. Wang, and Y. Ihara. Longer forms of amyloid protein: implications for the mechanism of intramembrane cleavage by *γ*-secretase. J. Neurosci., 25:436–445, 2005.

[22] M. Takami, Y. Nagashima, Y. Sano, S. Ishihara, M. Morishima-Kawashima, S. Funamoto, and Y. Ihara. *γ*-secretase: successive tripeptide and tetrapeptide release from the transmembrane domain of *β*-carboxyl terminal fragment. J. Neurosci., 29:13042–13052, 2009.

[23] M. Okochi, S. Tagami, K. Yanagida, M. Takami, T. S. Kodama, K. Mori, T. Nakayama, Y. Ihara, and M. Takeda. *γ*-secretase modulators and presenilin 1 mutants act differently on presenilin/*γ*-secretase function to cleave a*β*42 and *aβ*43. Cell Rep., 3:42–51, 2013.

[24] L. Sun, R. Zhou, G. Yang, and Y. Shi. Analysis of 138 pathogenic mutations in presenilin-on the in vitro production of a*β*_42_ and a*β*_40_ peptides in *γ*-secretase. Proc. Natl. Acad. Sci., 114:E476–485, 2016.

[25] B. I. P’erez-Revuelta, A. Fukumori, S. Lammich, A. Yamasaki, C. Haass, and H. Steiner. Requirement for small side chain residues within the gxgd-motif of presenilin for *γ* – secretase subsrate cleavage. J. Neurochem., 112:950–950, 2010.

[26] B. Kretner, A. Fukumori, P-H. Kuhn, B. I. Perez-Revuelta, S. F. Lichtenthaler, C. Haass, and H. Steiner. Imoprtant functional role of residue x of the presenilin gxgd protease active site motif for app substrate cleavage specificity and substrate selectivity of *γ* – secretase. J. Neurochem., 125:144–156, 2013.

[27] C. J. Crump, D. S. Johnson, and Y.-M. Li. Development and mechanism of *γ*-secretase modulators for alzheimer’s disease. Biochemistry, 52:3197–3216, 2013.

[28] A. Fukumori and H. Steiner. Substrate recruitment of *γ*-secretase and mechanism of clinical presenilin mutations revealed by photoaffinity mapping. EMBO J., e201694151, 2016.

[29] N. Watanabe, S. Takagi, A. Tominaga, T. Tomita, and T. Iwatsubo. Functional analysis of the transmembrane domains of presenilin 1 participation of transmembrane domains and 6 in the formation of initial substrate-binding site of *γ*-secretase. J. Biol. Chem., 285:19738–19746, 2010.

[30] M. Hitzenberger and M. Zacharias. *γ*-secretase studied by atomistic molecular dynamics simulations: Global dynamics, enzyme activation, water distribution and lipid binding. Front. Chem., 6:640, 2018.

[31] A. K. Somavarapu and K. P. Kepp. Membrane dynamics of *γ*-secretase provides a molecular basis for a*β* binding and processing. ACS Chem. Neurosci., 8:2424–2436, 2017.

[32] R. Kong, S. Chang, W. Xia, and S. T. C. Wong. Molecular dynamics simulation study reveals potentials substrate entry path into *γ*-secretase/presenilin-1. J. Struct. Biol., 191:120–129, 2015.

[33] A. Y. Kornilova, C. Das, and M. S. Wolfe. Differential effects of inhibitors on the *γ* – secretase complex. J. Biol. Chem., 278:16470–16473, 2003.

[34] C. Das, O. Berezovska, T. S. Diehl, C. Genet, I. Buldyrev, J-Y. Tsai, B. T. Hyman, and M. S. Wolfe. Designed helical peptides inhibit an intramembrane protease. J. Am. Chem. Soc., 125:11794–11795, 2003.

[35] F. Kamp, E. Winkler, J. Trambauer, A. Ebke, R. Fluhrer, and H. Steiner. Intramembrane proteolysis of *γ*-amyloid precursor protein by *γ*-secretase is an unusually slow process. Biophys. J., 108:1229–1237, 2015.

[36] R. Aguayo-Ortiz, C. Chavez-Garcia, J. E. Straub, and L. Dominguez. Characterizing the structural ensemble of *γ*-secretase using a multiscale molecular dynamics approach. Chem. Sci., 8:5576–5584, 2017.

[37] P. J. Barrett, Y. Song, W. D. Van Horn, E. J. Hustedt, J. M. Schafer, A. Hadziselimovic, A. J. Beel, and C. R. Sanders. The amyloid precursor protein has a flexible transmembrane domain and binds cholesterol. Science, 336:1168–1171, 2012.

[38] A. Goetz, N. Mylonas, P. Hoegel, M. Silber, H. Heinel, S. Menig, A. Vogel, H. Freyrer, D. Huster, B. Luy, D. Langosch, C. Scharnagl, C. Muhle-Groll, F. Kamp, and H. Steiner. Stabilization / destabilization of the app transmembrane domain by mutations in the diglycine hinge alter helical structure and dynamics, and impair cleavage by -secretase. bioRxiv, 10.1101/375006, 2018.

[39] O. Pester, P. J. Barrett, D. Hornburg, P. Hornburg, R. Pröbstle S. Widmaier, C. Kutzner, M. Dürrbaum, A. Kapurniotu, C. R. Sanders, C. Scharnagl, and D. Langosch. The backbone dynamics of the amyloid precursor protein transmembrane helix provides a rationale for the sequential cleavage mechanism of-secretase. J. Am. Chem. Soc., 135:1317–1329, 2013.

[40] L. Dominguez, S. C. Meredith, J. E. Straub, and D. Thirumalai. Transmembrane fragment structures of amyloid precursor protein depend on membrane surface curvature. J. Am. Chem. Soc., 136:854–857, 2014.

[41] C. Scharnagl, O. Pester, P. Hornburg, D. Hornburg, A. Gotz, and D. Langosch. Side-chain to main-chain hydrogen bonding controls the intrinsic backbone dynamics of the amyloid precursor protein transmembrane helix. Biophys. J., 106:1318–1326, 2014.

[42] J. Y. Lee, Z. Feng, X. Q. Xie, and I. Bahar. Allosteric modulation of intact *γ*-secretase structural dynamics. Biophys. J., 113:2634–2649, 2017.

[43] K. Yu, G. Yang, and J. Labahn. High-efficient production and biophysical characterisation of nicastrin and its interaction with appc100. Sci. Rep., 7:44297, 2017.

[44] T. L. Kukar, T. B. Ladd, P. Robertson, S. A. Pintchovski, B. Moore, M. A. Bann, Z. Ren, K. Jansen-West, K. Malphrus, S. Eggert, H. Maruyama, B. A. Contrell, P. Das, G. S. Basi, E. H. Koo, and T. E. Golde. Lysine 624 of the amyloid precursor protein (app) is a critical determinant of amyloid beta peptide length: Support for sequential model of *γ*-secretase intramembrane proteolysis and regulation by the app juxtamembrane region. J. Biol. Chem., 286:39804–39812, 2011.

[45] N. Clemente, A. Abdine, I. Ubarretxena-Belandia, and C. Wang. Coupled transmembrane substrate docking and helical unwinding in intramembrane proteolysis of amyloid precursor protein. Sci. Rep., 8:12411, 2018.

[46] H. Steiner, M. Kostka, H. Romig, G. Basset, B. Pesold, J. Hardy, A. Capell, L. Meyn, M. L. Grim, RF. Baumeister, K. Fechteler, and C. Haass. Glycine 384 is required for presenilin-1 function and is conserved in bacterial polytopic aspartyl proteases. Nat. Cell. Biol., 2(11):848, 2000.

[47] A. Y. Kornilova, F. Bihel, C. Das, and M. S. Wolfe. The initial substrate-binding site of *γ*-secretase is located on presenilin near the active site. Proc. Natl. Acad. Sci., 102:32303235, 2005.

[48] S. Padmanabhan, S. Marqusee, T. Ridgeway, T. T. Laue, and R. L. Baldwin. Relative helix-forming tendencies of nonpolar amino acids. Nature, 344(6263):268–270, 1990.

[49] K. T. O’Neil and W. F. DeGrado. A thermodynamic scale for the helix-forming tendencies of the commonly occurring amino acids. Science, 250(4981):646–651, 1990.

[50] P. C. Lyu, J. C. Sherman, A. Chen, and N. R. Kallenbach. α-helix stabilization by natural and unnatural amino acids with alkyl side chains. Proc. Natl. Acad. Sci., 88:5317–5320, 1991.

[51] S. Scheuermann, B. Hambsch, L. Hesse, J. Stumm, C. Schmidt, D. Beher, T. A. Bayer, K. Beyreuther, and G. Multhaup. Homodimerization of amyloid precursor protein and its implication in the amyloidogenic pathway of alzheimer’s disease. J. Biol. Chem., 276:33923–33929, 2001.

[52] L.-M. Munter, P. Voigt, A. Harmeier, D. Kaden, K. E. Gottschalk, C. Weise, R. Pipkorn, M. Schaefer, D. Langosch, and G. Multhaup. Gxxxg motifs within the amyloid precursor protein transmembrane sequence are critical for the etiology of a42. EMBO J., 26:17021712, 2007.

[53] E. Winkler, A. Julius, H. Steiner, and D. Langosch. Homodimerization protects the amyloid precursor protein c99 fragment from cleavage by -secretase. Biochemistry, 54:6149–6152, 2015.

[54] Y. Yan, T-H. Xu, K. Melcher, and H. E. Xu. Defining the minimum substrate and charge recognition model of gamma-secretase. Acta Pharm. Sinic., 38:1412–1424, 2017.

[55] M. A. Lomize, I. Pogozheva, H. Joo, H. I. Mosberg, and A. L. Lomize. Opm database and ppm web server: resources for positioning of proteins in membranes. Nucleic Acids Res., 40:D370–6, 2012.

[56] S. Jo, V. G. Iyer, and W. Im. Charmm-gui: A web-based graphical user interface for charmm. J. Comput. Chem., 29:1859–1865, 2008.

[57] E. L. Wu, X. Cheng, S. Jo, H. Rui, K. C. Song, E. M. Dvila-Contreras, Y. Qi, J. Lee, V. Monje-Galvan, R. M. Venable, J. B. Klauda, and W. Im. Charmm-gui membrane builder toward realistic biological membrane simulations. J. Comput. Chem., 35:1997–2004, 2014.

[58] R. Singh, A. Barman, and R. Prabhakar. Computational insights into aspartyl protease activity of presenilin 1 (ps1) generating alzheimer amyloid *β*-peptides (a*β*_40_ and a*β*_42_). J. Phys. Chem. B., 113:2990–2999, 2008.

[59] J. Nickolls, I. Buck, M. Garland, and K. Skadron. Scalable parallel programming with cuda. ACM Queue, 6(2):40–53, 2008.

[60] A. W. Goetz, M. J. Williamson, D. Xu, D. Poole, S. Le Grand, and R. C. Walker. Routine microsecond molecular dynamics simulations with amber – part i: Generalized born. J. Chem. Theory Comput., 8:1542–1555, 2012.

[61] R. Salomon-Ferrer, A. W. Goetz, D. Poole, S. Le Grand, and R. C. Walker. Routine microsecond molecular dynamics simulations with amber – part ii: Particle mesh ewald. J. Chem. Theory Comput., 9:3878–3888, 2013.

[62] S. Le Grand, A. W. Goetz, and R. C. Walker. Spfp: Speed without compromise – a mixed precision model for gpu accelerated molecular dynamics simulations. Comp. Phys. Comm., 184:374–380, 2013.

[63] D. A. Case, R. M. Betz, D. S. Cerutti, III T. E. Cheatham, T. A. Darden, R. E. Duke, T. J. Giese, A. W. Goetz H. Gohlke, N. Homeyer, S. Izadi, P. Janowski, J. Kaus, A. Kovalenko, T. S. Lee, S. LeGrand, P. Li, C. Lin, T. Luchko, R. Luo, B. Madej, D. Mermelstein, K. M. Merz, G. Monard, H. Nguyen, H. T. Nguyen, I. Omelyan, A. Onufriev, D. R. Roe, A. Roitberg, C. Sagui, C. L. Simmerling, W. M. Botello-Smith, J. Swails, R. C. Walker, J. Wang, R. M. Wolf, X. Wu, L. Xiao, and P. A. Kollman. Amber 2016. University of California, San Francisco, 2016.

[64] J. A. Maier, C. Martinez, K. Kasavajhala, L. Wickstrom, K. Hauser, and C. Simmerling. ff14sb: improving the accuracy of protein side chain and backbone parameters from ff99sb. J. Chem. Theory Comput., 11(8):3696–3713, 2015.

[65] C. J. Dickson, D. M. Madej, Å. A. Skjevik, R. M. Betz, K. Teigen, I. R. Gould, and R. C. Walker. Lipid14: The amber lipid force field. J. Chem. Theory Comput., 10:865–879, 2014.

[66] P. Mark and L. Nilsson. Structure and dynamics of the tip3p, spc, and spc/e water models at 298 k.. J.Phys. Chem. A, 105(43):9954–9960, 2001.

[67] T. Darden, D. York, and L. Pedersen. Particle mesh ewald: An n log (n) method for ewald sums in large systems. J. Chem. Phys., 98(12):10089–10092, 1993.

[68] N. Goga, A. J. Rzepiela, A. H. De Vries, S. J. Marrink, and H. J. C. Berendsen. Efficient algorithms for langevin and dpd dynamics. J. Chem. Theory Comput., 8:3637–3649, 2012.

[69] J. Aqvist, P. Wennerstrom, M. Nervall, S. Bjelic, and B. O. Brandsdal. Molecular dynamics simulations of water and biomolecules with a monte carlo constant pressure algorithm. Chem. Phys. Lett., 384:288–294, 2004.

[70] J.P. Ryckaert, G. Ciccotti, and H.J.C. Berendsen. Numerical integration of the cartesian equations of motion of a system with constraints: molecular dynamics of n-alkanes. J. Comput. Phys., 23(3):2327–2341, 1977.

[71] C. W. Hopkins, S. Le Grand, R. C. Walker, and A. E. Roitberg. Long-time-step molecular dynamics through hydrogen mass repartitioning. J. Chem. Theory Comput., 11:1864–1874, 2015.

[72] D. R. Roe and T. E. Cheatham III. Ptraj and cpptraj: Software for processing and analysis of molecular dynamics trajectory data. J. Chem. Theory Comput., 9:3084–3095, 2013.

[73] W. Humphrey, A. Dalke, and K. Schulten. Vmd – visual molecular dynamics. J. Mol. Graphics, 14:33–38, 1996.

[74] S. Genheden and U. Ryde. The mm/pbsa and mm/gbsa methods to estimate ligand-binding affinities. Expert Opin. Drug Discov., 10:449–461, 2015.

[75] B. R. Miller III, T. D. McGee Jr., J. M. Swails, N. Homeyer, H. Gohlke, and A. E. Roitberg. Mmpbsa.py: An efficient program for end-state free energy calculations. J. Chem. Theory Comput., 8:3314–3321, 2012.

[76] F. Fogolari, A. Brigo, and H. Molinari. The poisson-boltzmann equation for biomolecular electrostatics: a tool for structural biology. J. Mol. Recognit., 15:377–392, 2002.

