## Supplemental Information for "Structural Modeling of *γ*-Secretase A*β_n_* Complex Formation and Substrate Processing"

### Supporting Information: Structural Modeling of $\gamma$-Secretase A$\beta_n$ Complex Formation and Substrate Processing

*M. Hitzenberger and M. Zacharias*

Physics Department T38

Technical University of Munich

James-Frank-Str. 1

85748 Garching, Germany

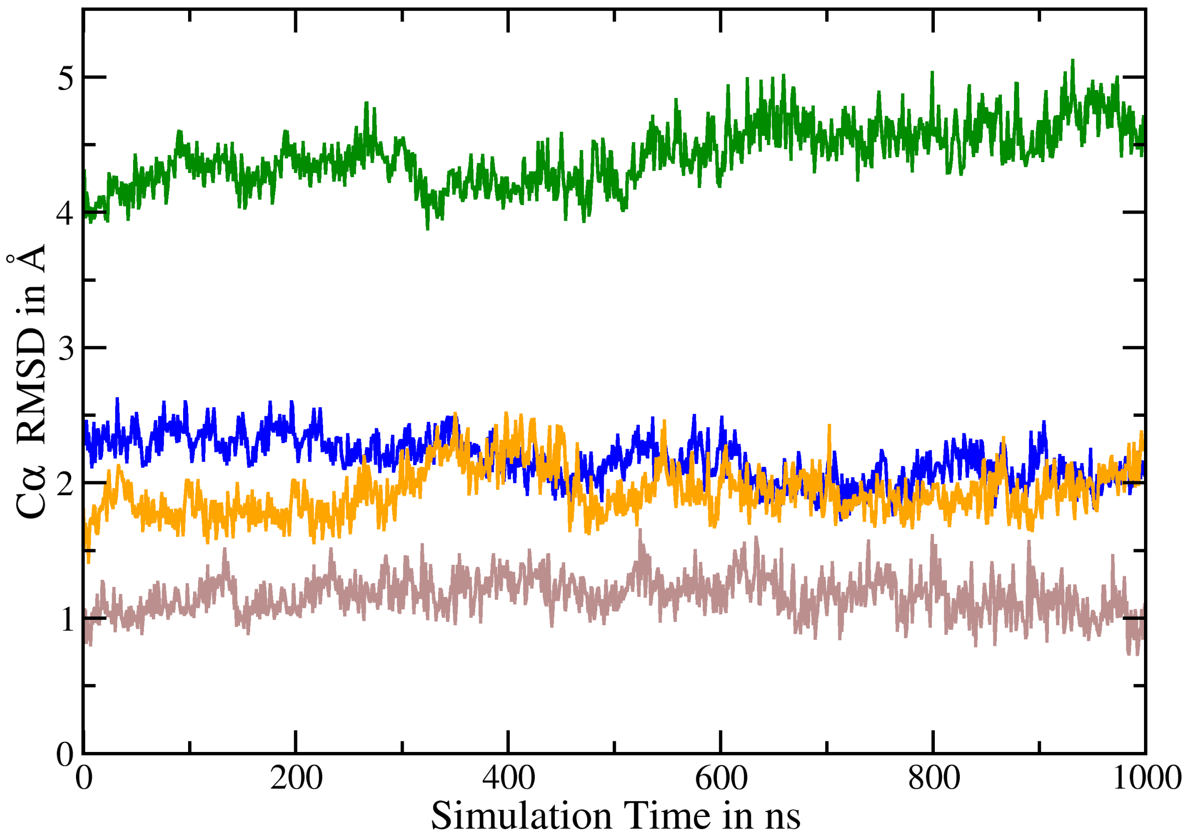


**Figure 1:** C$\alpha$ RMSD of the apo-state simulation with PDB:5FN2 as the reference structure. Green: Nicastrin; blue: TMDs of PS-1; orange: TMDs of Aph-1a, brown: TMDs of Pen-2.

| | Steps | Time Step | K Proteins | K Lipids | Temperature | Pressure |
|---|---|---|---|---|---|---|
| **min** | 50000 | minimization | 10.0 | 2.5 | — | — |
| **eq 1** | 25000 | 1ps | 10.0 | 2.5 | 0K to 303.15K | NVT |
| **eq 2** | 25000 | 1ps | 5.0 | 2.5 | 303.15K | NVT |
| **eq 3** | 25000 | 1ps | 2.5 | 1.0 | 303.15K | 0bar to 1bar |
| **eq 4** | 50000 | 2ps | 1.0 | 0.5 | 303.15K | 1bar |
| **eq 5** | 50000 | 2ps | 0.5 | 0.1 | 303.15K | 1bar |
| **eq 6** | 50000 | 2ps | 0.1 | 0.0 | 303.15K | 1bar |

**Table 1:** Overview over the 7 pre-equilibration steps, performed prior to the actual 500ns equilibration. Positional restraints, K are given in kcal/mol*Å$^2$ (restraints are in AMBER format: K*$\Delta x^2$)

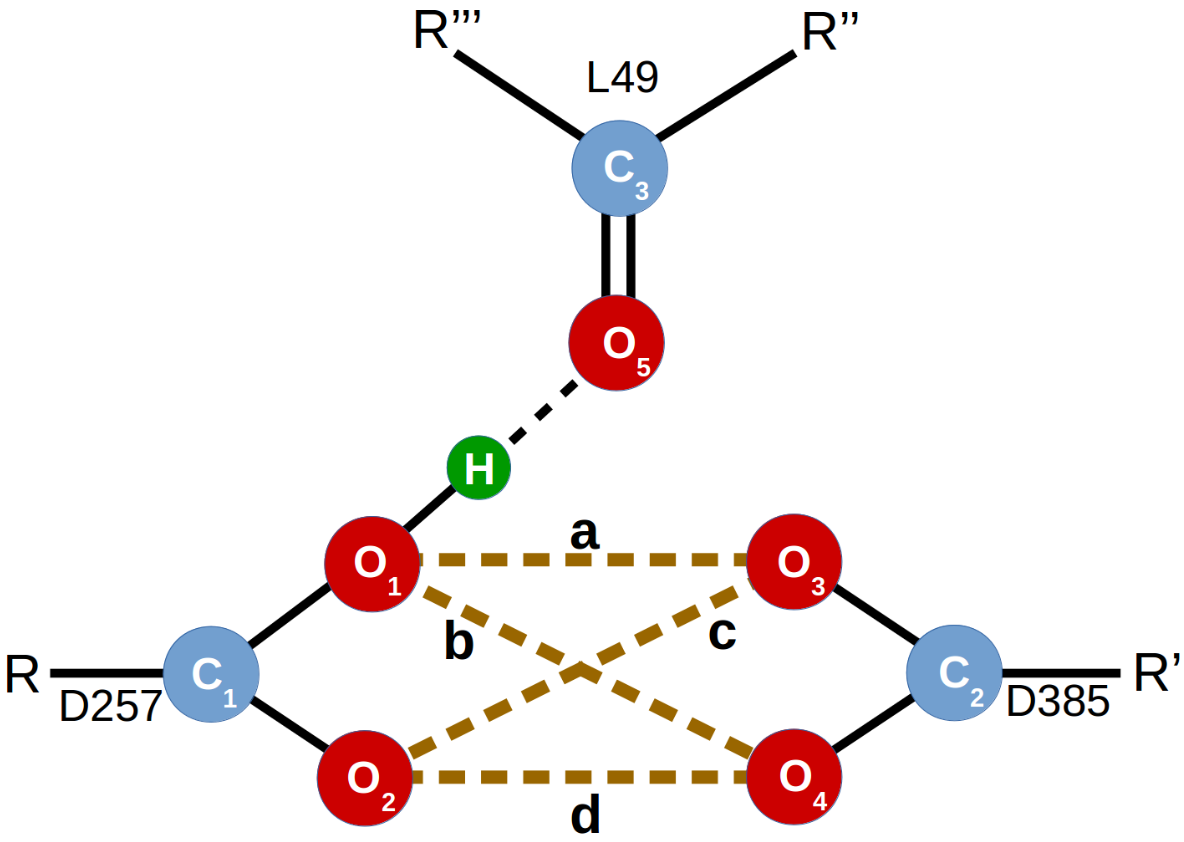


**Figure 2:** Schematic of the desired active site-L49 coordination in simulation GC99.

The positions of the energy minima of the aspartate distance restraints were taken from experimental active site geometries of aspartyl proteases.[1] The geometry can be taken from Figure 2. The desired distances were: a=3.0Å, b=4.5Å, c=4.8Å, d=5.8Å and 1.8Å for the hydrogen bond between H and O5. Additionally, a center of mass restraint between atoms C1, C2. O1, O2, O3, O4 (active site) and O5 (substrate) with a desired distance of 2.0Å was used to keep the active site-substrate interface in a compact conformation for the entirety of the sampling time. The force constants were 5kcal/mol*Å$^2$ in all cases (restraints are in AMBER format: K*$\Delta x^2$).

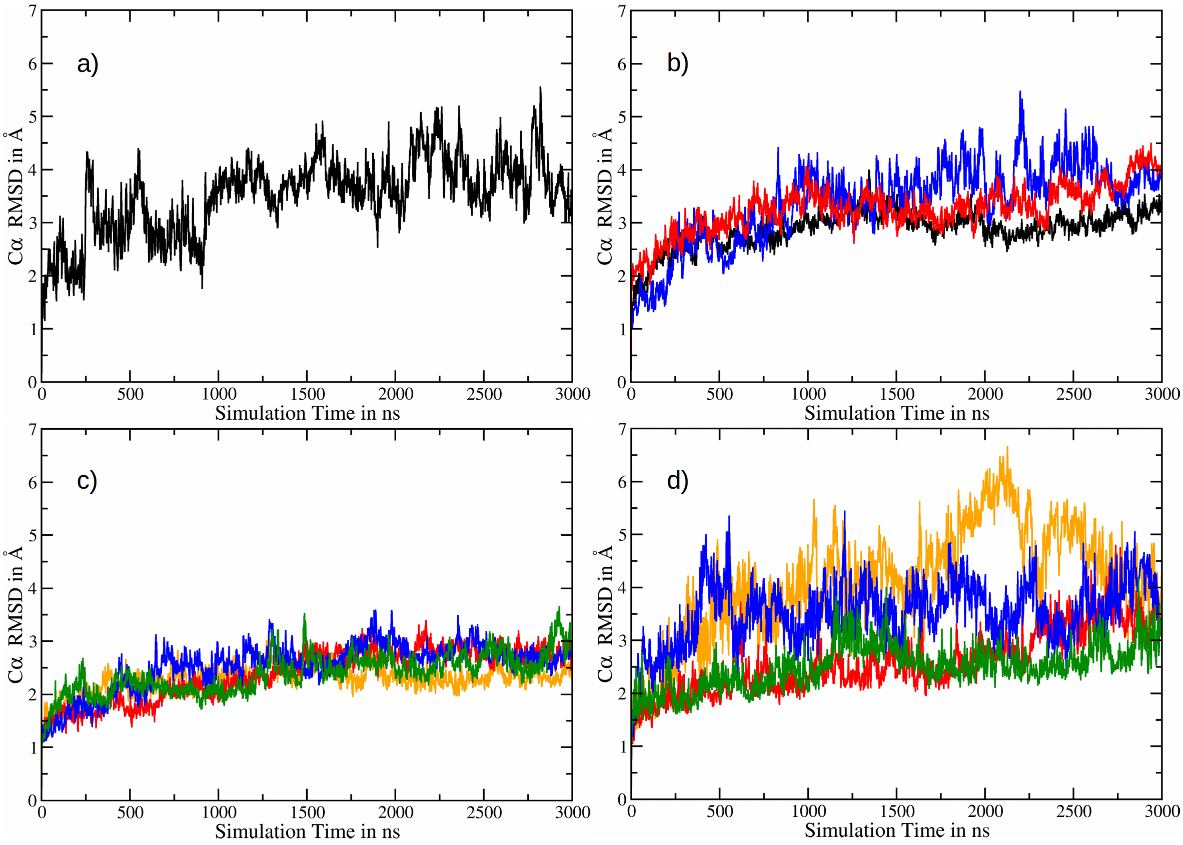


**Figure 3:** C$\alpha$ RMSDs of simulations a) C99, b) Model 1 (black), Model 2 (blue) and Model 3 (red). c) GC99* (orange), GA$\beta_{49}$* (red), GA$\beta_{46}$* (blue) and GA$\beta_{43}$* (green). d) GC99 (orange), GA$\beta_{49}$ (red), GA$\beta_{46}$ (blue), GA$\beta_{43}$ (green).
In case of simulations including GSEC, the very mobile loop region between residues 263 to 383 has been left out from the RMSD calculations.

| Residue | FAD Mutation(s) | Interaction Site(s) |
|---|---|---|
| K16 | N | nicastrin |
| E22 | G | nicastrin |
| D23 | N | nicastrin |
| L34 | V | — |
| A42 | T | PS-TMD 3 |
| T43 | A, I | PS-TMDs 5, 7 |
| V44 | A, M | PS-TMD 2 |
| I45 | F, M, T, V | PS-TMDs 2, 3 |
| V46 | F, G, I, L | PS-TMDs 3, 5; GXGD |
| T48 | P | PS-TMDs 2, 6 |
| L52 | P | GXGD, loop 6* |
| K53 | N | PS-TMD 2 |
| L49 | W †, Y † | GXGD |

**Table 2:** Overview over the GSEC binding sites of known C99 FAD - causing mutation sites (with the exception of L49).
* loop 6 denotes the large intracellular loop region between TMDs 6 and 7 (which is cleaved upon maturation).
† Mutations that are not known to cause FAD but impede $\epsilon$-cleavage.[2]

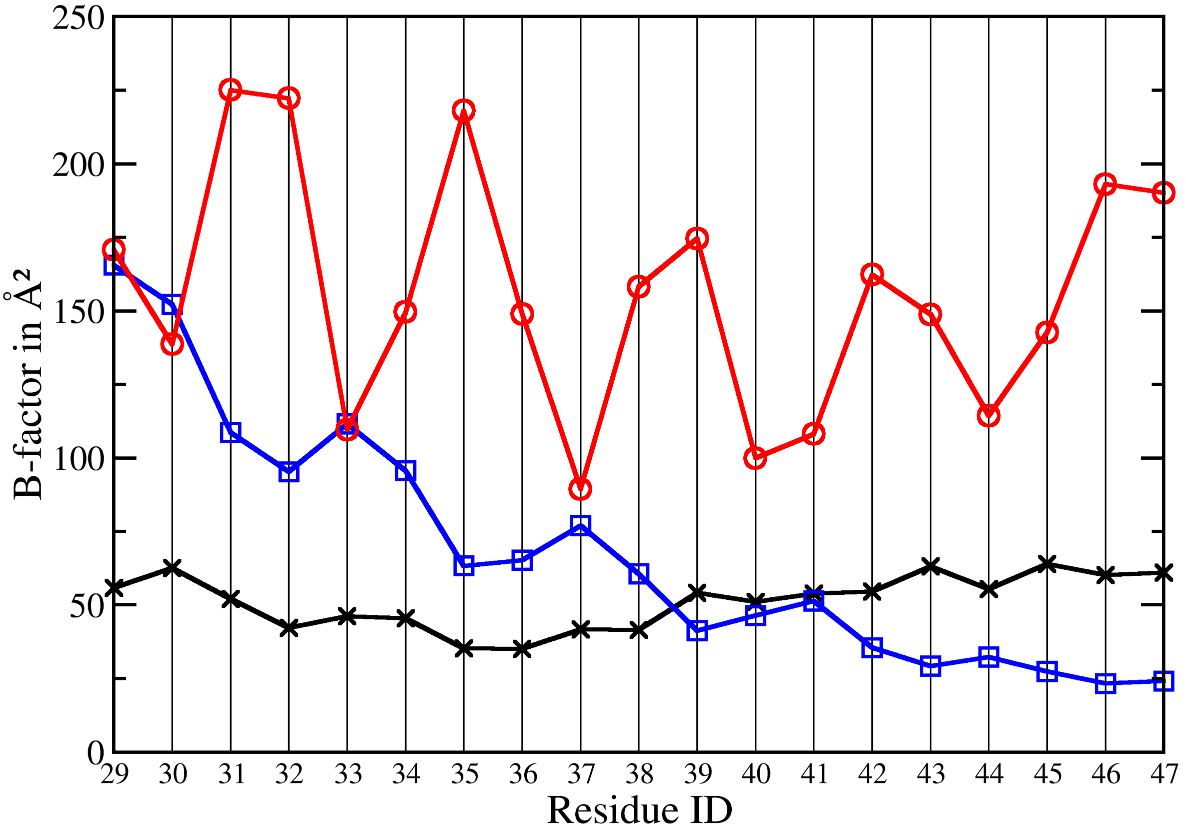


**Figure 4:** B-factor calculations of the C99 helix in models 1 (black), 2 (blue) and 3 (red). Only the C$\alpha$ atoms have been considered.

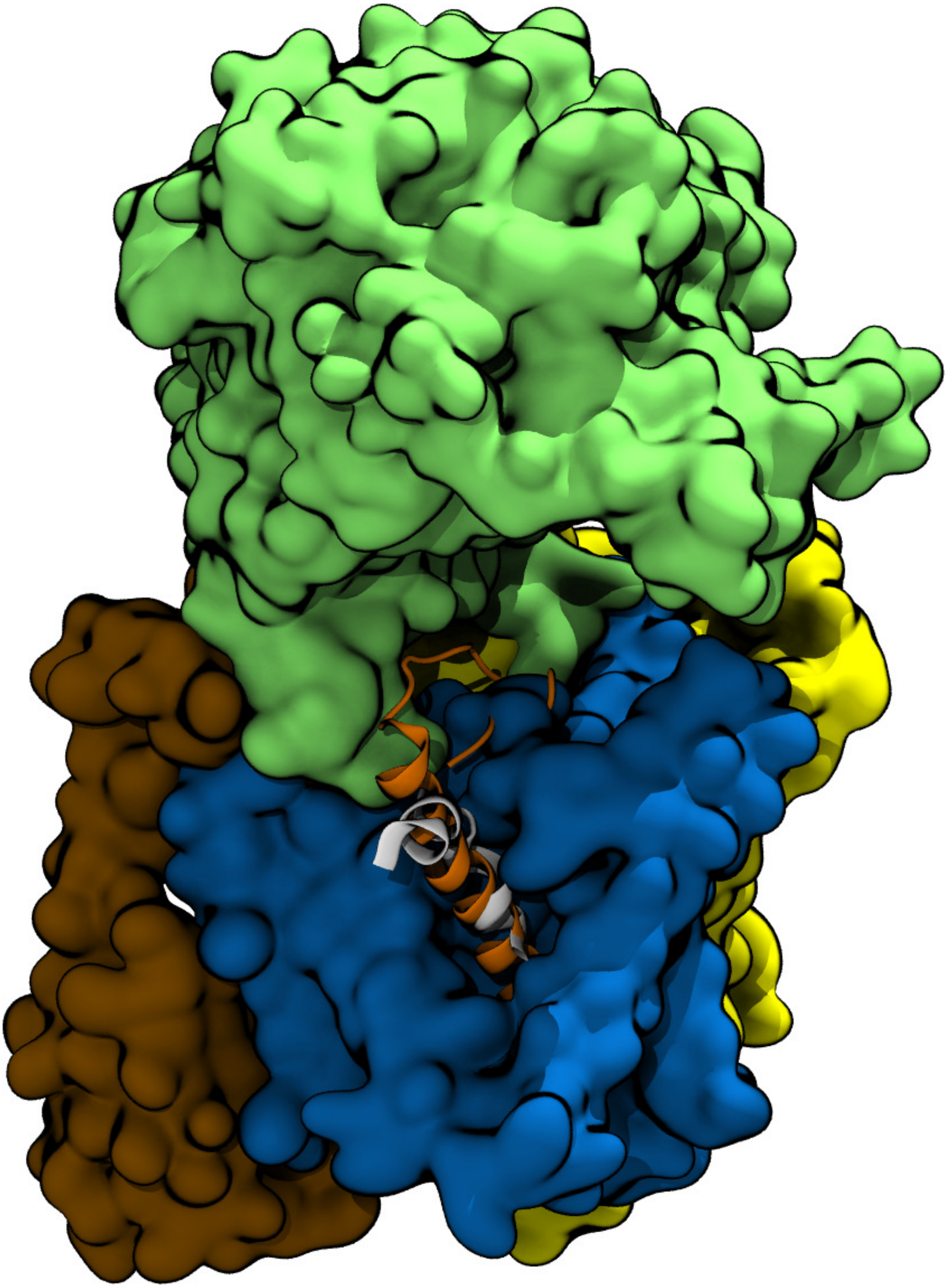

**Figure 5:** Superposition of simulation GC99 and PDB structure 5FN3, highlighting that the positions of the co-purifying helix (white) and C99 (orange) in our model, coincide. The GSEC structure is taken from the PDB file.

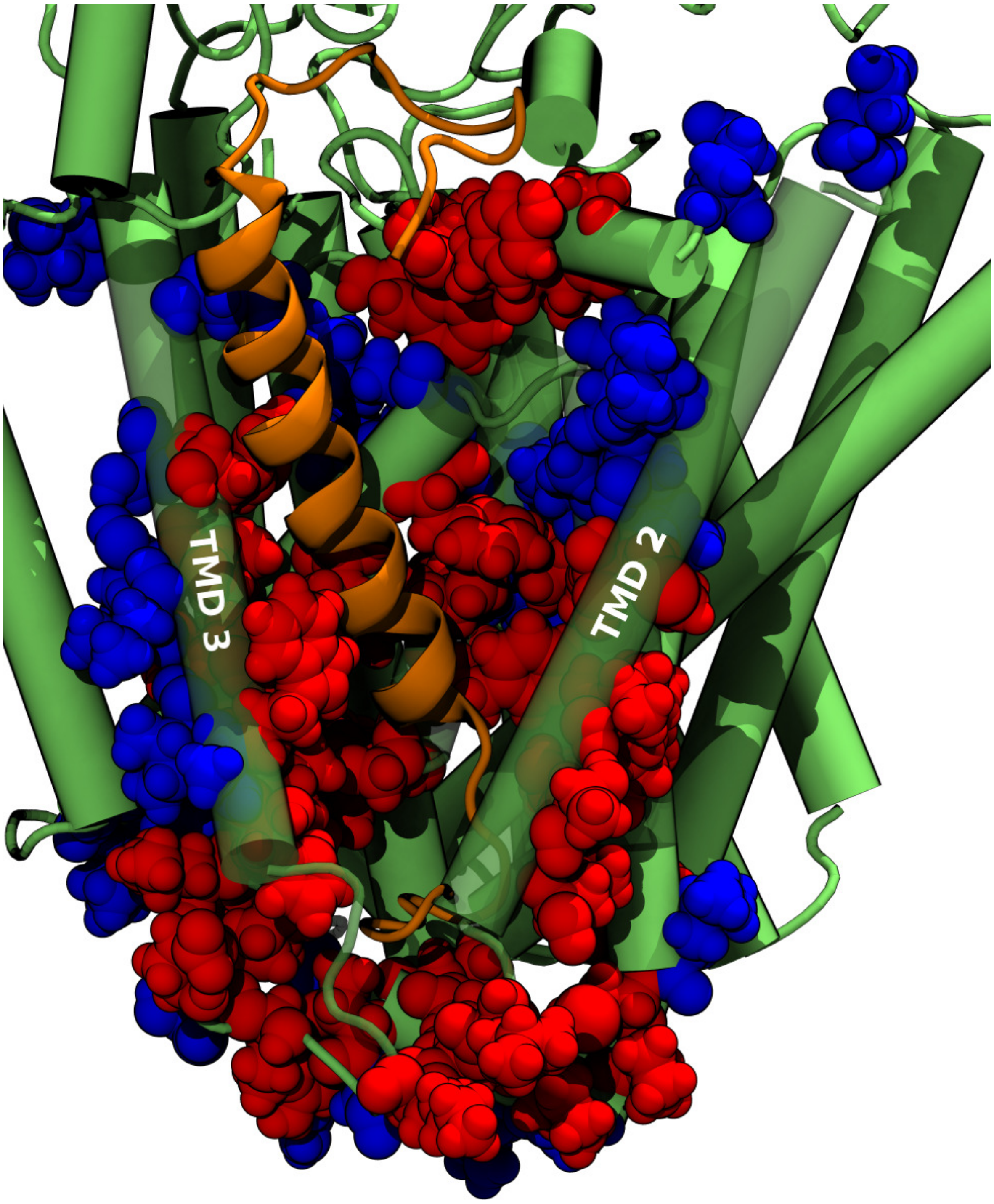


**Figure 6:** Snapshot from simulation GC99, depicting the binding interface of the E-S complex. C99 (orange) is coordinated by GSEC (green). The residues highlighted in red are known FAD-mutation sites, within 5Å of C99. Blue residues are mutation sites that are more separated from C99.

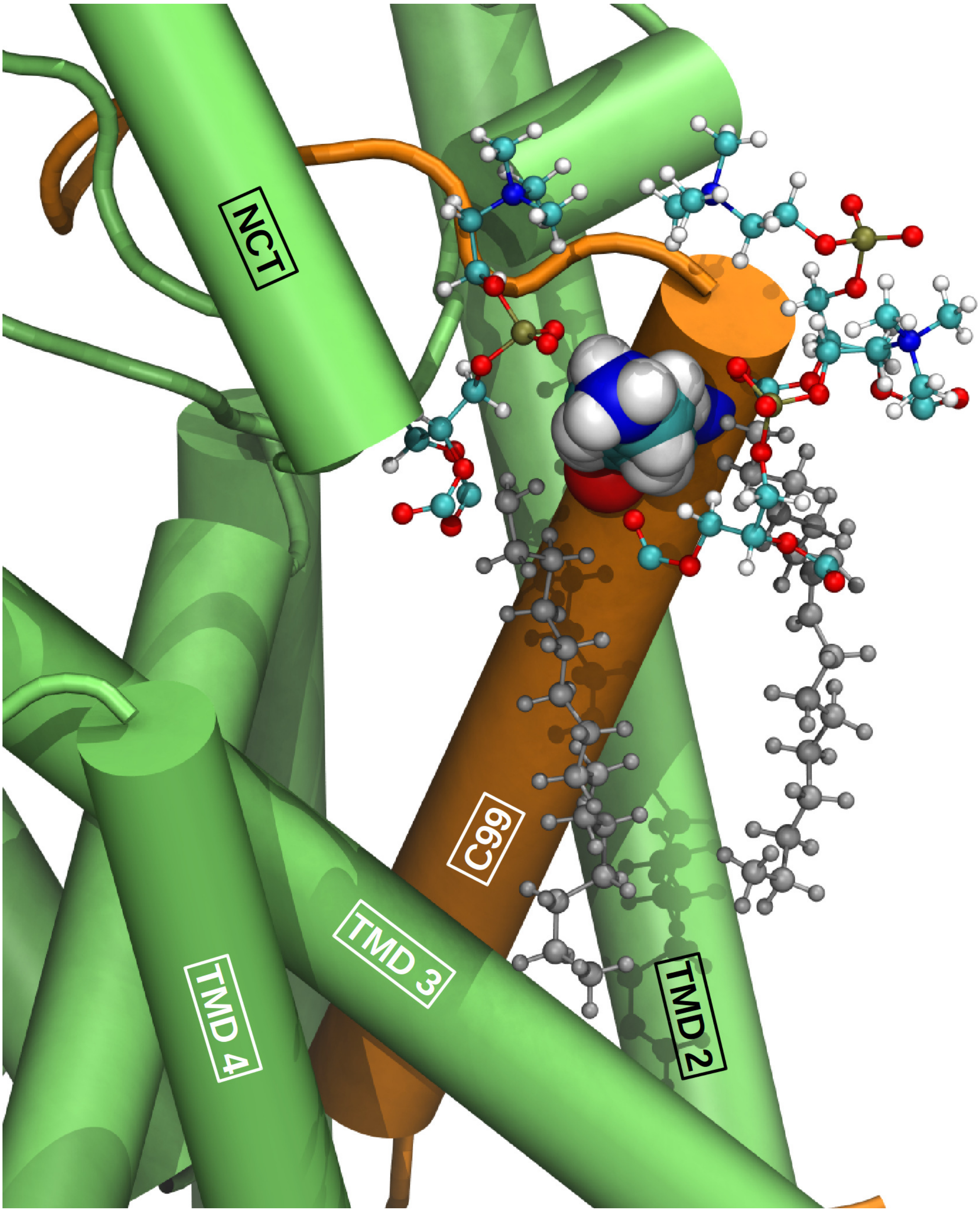


**Figure 7:** Snapshot from simulation GC99, depicting the characteristic interaction between K28 (shown as Van-der-Waals spheres) and two POPC molecules (shown in ball and stick representation).

#### References

[1] R. Singh, A. Barman, and R. Prabhakar. Computational insights into aspartyl protease activity of presenilin 1 (ps1) generating alzheimer amyloid $\beta$-peptides (a$\beta$40 and a$\beta$42). *J. Phys. Chem. B.*, 113:2990–2999, 2008.

[2] T. Xu, Y. Yan, Y. Kang, Y. Jiang, K . Melcher, and H. E. Xu. Alzheimer's disease-associated mutations increase amyloid precursor protein resistance to $\gamma$-secretase cleavage and the a$\beta$40/a$\beta$42 ratio. *Cell Discovery*, 1:16026, 2016.
